# Modeling VEGF and GLUT1 Expression as Coadapted Foraging Strategies in Cancer

**DOI:** 10.64898/2026.04.14.718570

**Authors:** Ranjini Bhattacharya, Robert A. Gatenby, Joel S. Brown

## Abstract

Natural selection acting on cancer cells within their tumor microenvironment should favor cells with fast or efficient nutrient uptake strategies. Here, we develop and analyze a game-theoretic model focusing on the coadaptation between two foraging traits: vascular endothelial growth factor (VEGF) and glucose transporter 1 (GLUT1). Studies show that VEGF and GLUT1 are often co-expressed and are associated with more aggressive tumor phenotypes and poor clinical prognosis. VEGF is a diffusible paracrine factor that recruits blood vessels towards neighborhoods of cancer cells (angiogenesis). GLUT1 is a cell-surface transporter that enables the uptake of nutrients, especially glucose. We model these strategies operating at different scales: VEGF influences resource availability at the neighborhood level, while GLUT1 determines resource uptake at the cellular level. For VEGF, we introduce a resource-sharing continuum. With no resource sharing, cells access resources in proportion to their VEGF contribution. With uniform sharing, cells have equal access to resources, regardless of their VEGF contribution. The former leads to a tragedy of the commons and overproduction of VEGF. The latter yields a public goods game with moderate VEGF expression matching a group optimum. GLUT1 expression mediates uptake of resources recruited by VEGF and is largely independent of the degree of resource sharing. Therapeutically, both VEGF and GLUT1 inhibitors are more effective in high resource-sharing neighborhoods and less so as resource sharing declines. Overall, inhibition of GLUT1-mediated uptake emerges as more effective. The model, perhaps the first to consider VEGF and GLUT1 as coadaptations, emphasizes the need to consider cancer cell traits jointly.

## 1. Introduction

Cancer cells exist in a resource-limited tumor microenvironment. This scarcity arises from both inadequate vascular supply and rapid nutrient consumption by proliferating cells [1,2]. In response, cancer cells modulate a range of signaling pathways and transport mechanisms, including the expression of vascular endothelial growth factor (VEGF), which promotes angiogenesis, and glucose transporter 1 (GLUT1), which facilitates glucose uptake [3–6]. While VEGF expression increases blood supply within a region, GLUT1 expression increases a cell’s capacity to take up glucose. Together, these responses act in concert as a coadaptation [7,8] for optimizing resource acquisition in cancer cells. Evolutionary pressures within the tumor should shape the cell’s investment in these two strategies and determine how expression of one influences the expression of the other. In this study, we will explore how these complementary strategies evolve together under shared constraints.

In response to hypoxia and resource deprivation in the tumor, the master transcriptional regulator, hypoxia inducible factor-1 (HIF-1), is activated [9,10]. Among its downstream targets are VEGF and GLUT1. VEGF is a paracrine signaling factor released into the tumor microenvironment that instigates the angiogenic cascade. It binds to VEGF receptors (primarily VEGFR-2) on endothelial cells lining existing blood vessels, activating signaling pathways that promote endothelial cell proliferation, migration, and sprouting toward regions of high VEGF concentration [11,12]. The local concentration of VEGF in a region influences the extent of neovascular growth. In addition to its paracrine effects, VEGF can also bind to VEGF receptors expressed on cancer cells, promoting autocrine signaling that aids cellular proliferation [13,14].

Glucose transporters (GLUTs) belong to the SLC2A family of facilitative membrane transporters and mediate glucose uptake [6]. Among the fourteen isoforms, GLUT1 is the most consistently upregulated in cancers and is used clinically as the basis for fluorodeoxyglucose positron emission tomography (FDG-PET) imaging [5,15–18]. GLUT1 localizes to the plasma membrane and increases cellular glucose influx, supporting the elevated glycolytic demands of cancer cells. Other isoforms, such as GLUT3, are also upregulated in tumors with high metabolic demand [19–21]. Although glucose remains the primary substrate driving tumor metabolism, certain GLUT isoforms can transport other hexoses and related molecules, such as dehydroascorbic acid, reflecting the metabolic plasticity of cancer cells under resource-limited conditions [22,23].

Co-expression of VEGF and GLUT1 is frequently associated with increased microvessel density, elevated metabolic activity, and more aggressive tumor phenotypes [24–26]. For example, studies in esophageal and ovarian cancers have reported associations between GLUT1 expression, VEGF expression, and measures of vascularization and glucose uptake [27]. In esophageal squamous cell carcinoma, GLUT1 expression has been shown to correlate with increased VEGF expression and FDG uptake [28]. These findings suggest a close relationship between these two traits.

These markers have also been evaluated for their prognostic significance. Elevated expression of VEGF and GLUT1, either individually or in combination with other markers such as carbonic anhydrase IX (CAIX), has been associated with poor survival, increased tumor aggressiveness, and resistance to therapy across multiple cancer types [29–32]. In some cases, a combined assessment of VEGF and GLUT1 provides greater prognostic value than the individual markers [31]. Together, these observations highlight the importance of considering these two adaptations as co-evolved strategies, rather than in isolation.

Tumor angiogenesis has been extensively modeled. Early continuum and hybrid models developed by Anderson and Chaplain [33,34], as well as Byrne and collaborators [35,36] laid the foundation for describing endothelial cell migration, chemotaxis, and vascular sprouting in response to angiogenic factors. Subsequent work has incorporated increasingly detailed mechanistic descriptions of VEGF signaling and transport. For example, Finley et al. developed compartmental models of VEGF distribution that account for VEGF isoforms, receptor binding, and interactions across tumor compartments [37,38]. Hybrid models such as those by Phillips et al. [39] and Nikmaneshi et al. [40] have coupled tumor growth with angiogenesis through feedback between hypoxia, VEGF production, and vascular supply. In addition to mechanistic and multiscale models, evolutionary and game-theoretic approaches have examined angiogenesis as a collective behavior, framing VEGF production as a public goods game, where individual cells balance the costs of VEGF production against shared benefits from VEGF expression [41,42].

In contrast to the extensive modeling of angiogenesis, mathematical models of tumor metabolism have largely focused on glucose as a diffusible resource, with less attention given to the regulatory mechanisms governing glucose uptake, such as GLUT1 expression. Early mathematical models of tumor growth, such as those developed by Greenspan [43], incorporated nutrient limitation through diffusion-based frameworks to describe how the availability of substrates such as oxygen and glucose constrain tumor expansion. Subsequent work by Casciari et al. [44] and Ward and King [45] extended these approaches by modeling the kinetics of glucose and oxygen consumption using reaction–diffusion equations, linking nutrient transport to cell proliferation, quiescence, and necrosis. Gatenby et al. [46] modeled glucose metabolism and acidosis using cellular automaton frameworks, to examine how gradients of oxygen, glucose, and acidity drive the selection of glycolytic phenotypes, providing a theoretical basis for the emergence of the Warburg effect in cancer [47–49]. Subsequent work has treated glycolytic and oxidative cells as distinct subpopulations. For example, Kareva et al. [50] modeled tumor–immune–glucose interactions using predator–prey dynamics. More recent approaches have incorporated glucose uptake into models of fluorodeoxyglucose (FDG) transport and uptake used to interpret PET imaging data [51,52]. Mechanistic models have also examined the role of glucose transport in tumor proliferation, including work by Ampatzoglou et al. [53], which explicitly incorporates GLUT1 expression into a multiscale tumor growth model.

Despite extensive empirical and theoretical work on angiogenesis and glucose metabolism, existing models have largely treated VEGF and GLUT1 expression as independent processes. To our knowledge, no modeling framework has examined these traits as coordinated, co-adapted strategies. Here, we address this gap.

In our earlier work, we developed an evolutionary game-theoretic model to study how cancer cells evolve VEGF expression as a function of local resource sharing [21]. We showed that in tumor neighborhoods with high resource sharing, where resources are distributed uniformly among cells, cells engage in a cooperative public goods game. In contrast, when resource sharing is limited, cells are incentivized to increase VEGF expression to secure a larger share of resources, giving rise to a tragedy of the commons [54,55]. In this regime, excessive individual investment in VEGF ultimately reduces overall tumor fitness. These results demonstrated the influence of cell-cell interactions on the evolution of VEGF as a foraging strategy.

Here, we build on this perspective and study the coadaptation of VEGF and GLUT1 expression as foraging strategies [56]. VEGF acts as a diffusible, paracrine signal that modulates resource availability at the neighborhood level, such that a cell’s access to nutrients depends on both its own expression and that of its neighbors [21]. In contrast, GLUT1 functions at the level of individual cells, with glucose uptake determined primarily by a cell’s own expression. These traits therefore, operate at different scales and impose distinct costs [57,58]: VEGF production requires sustained biosynthetic investment and generates shared benefits, whereas GLUT1 overexpression increases metabolic demand and glycolytic flux. Natural selection within the tumor microenvironment acts on the expression of these traits, shaping how cells allocate investment between these strategies under varying ecological and resource-sharing conditions.

Here, we model the “game” that emerges as cancer cells allocate investment to these two complementary foraging strategies within a tumor. Specifically, we 1) develop an evolutionary game-theoretic framework [59,60] in which the VEGF and GLUT1 are coadapted traits, 2) analyze how investment in these traits varies along a continuum of resource-sharing conditions, 3) analyze the effects of key parameters, and 4) explore the implications of these dynamics for therapeutic strategies targeting VEGF and GLUT1.

## 2. Methods

To investigate how cancer cells allocate investment between VEGF and GLUT1 expression, we consider a population of cancer cells organized into local neighborhoods within the tumor microenvironment. The neighborhood is described by cells that directly interact and influence each other’s access to nutrients supplied by shared blood vasculature [21,61]. The collective VEGF production of the neighborhood diverts blood flow towards it, determining the total resource supply. A part of this supply is available to each of the cancer cells in the neighborhood [62]. For a cancer cell to obtain its share, it must invest in GLUT1, which enables resource uptake from the blood. We model VEGF expression and GLUT1 expression as continuous quantitative traits representing investment in resource supply and uptake, respectively. VEGF acts at the neighborhood level by modulating resource availability, whereas GLUT1 acts at the cellular level by determining a cell’s ability to capture available resources.

We formalize these interactions using a fitness-generating function (*G*-function) [63–65], which describes the per capita growth rate of a focal cell within its neighborhood. A focal cell’s fitness is influenced by the number of cancer cells within its neighborhood (*N*), the average VEGF expression of the other *N* − 1 cells of the neighborhood (*u*), the focal cell’s GLUT1 expression (*y*), and the focal cell’s VEGF expression (*v*): *G* (*N, u, v, y*).

We assume that the total VEGF production of all cells, *x* = *v* + *u* (*N* − 1), influences the overall vasculature and resource supply to the neighborhood, and that both *v* and *u* can influence how much of this supply is available to the focal cell. Given a share of the resource supply, it is only the focal individual’s investment in GLUT1, *y*, and not that of others, which influences the uptake of its share. We develop these assumptions into the following model:

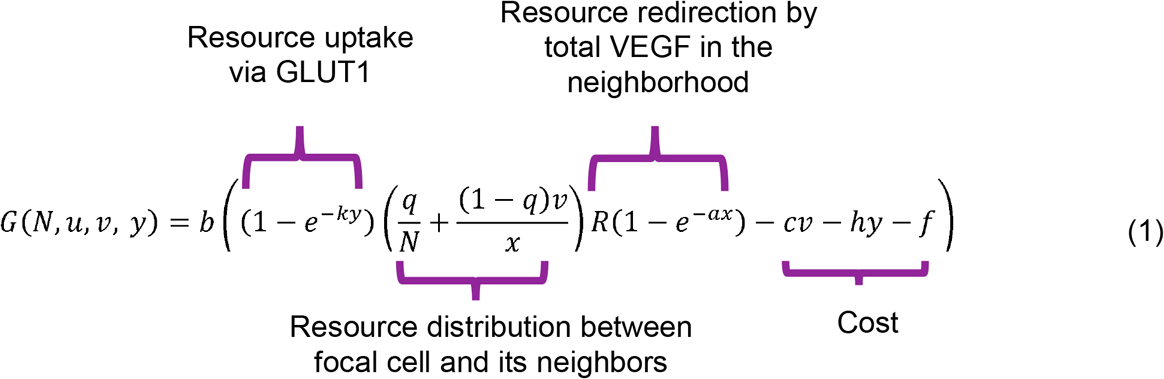

where *R* is the maximum possible resource influx, *a* is the efficacy of VEGF mediated resource redirection to the neighborhood, *k* is the efficacy of GLUT1 mediated resource uptake, *c* and *h* are the per unit costs of VEGF and GLUT1 expression, respectively, and *f* is the fixed cost of survival. In Eq. (1), *R* (1 − *e* ^−*ax*^ ) determines the total amount of resources entering a neighborhood as a function of *x*, the neighborhood’s total VEGF expression. *q* ∈ [0,1] is the degree of resource sharing in the neighborhood. When the degree of resource sharing is maximum (*q* = 1), resources are evenly distributed regardless of each cell’s VEGF contribution. When there is no resource sharing (*q* = 0), resources are allocated in proportion to each cell’s VEGF contribution. The term, (1 − *e* ^−*ky*^ ), represents GLUT1 mediated resource import, with the assumption that glucose uptake increases with GLUT1 expression but eventually saturates (as the function approaches 1).

The *G*-function models both cancer population dynamics and trait dynamics. The rate of evolution of the two traits, VEGF and GLUT1 expression, is a function of the selection gradient and the cell’s phenotypic plasticity in these traits. The cancer population (*N*), VEGF expression (*u*), and GLUT1 expression (*y*) dynamics are governed by Eq. (2), (3), (4).

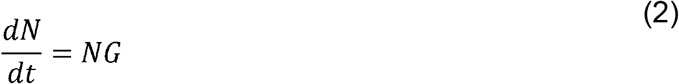

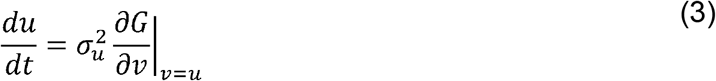

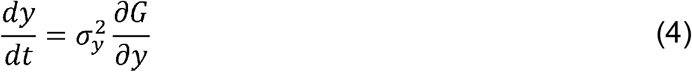

where 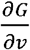 and 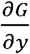 are the selection gradients with respect to the focal cell’s VEGF expression and GLUT1 expression respectively. The parameters 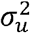 and 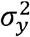 are the speed of phenotypic plasticity in VEGF and GLUT1 expression, respectively, in response to the strength of selection [23,66].

### 2.1 Modeling Therapy

To model therapy, we consider two drug modalities. The first are VEGF inhibitors or neutralizing agents that reduce the effectiveness of VEGF-mediated resource recruitment to the tumor (*a*). This class includes clinically approved drugs such as bevacizumab and sunitinib, which target VEGF signaling and angiogenesis [67–69]. The second are GLUT1 inhibitors, which reduce the efficacy of GLUT1-mediated glucose import into cells (*k*). While fewer GLUT1 inhibitors are currently in clinical use, several compounds are under development or study, including STF-31, WZB117, BAY-876, and ritonavir, which act through competitive or noncompetitive mechanisms to block glucose uptake [70–73]. We modify Eq. (1) to incorporate these therapies as follows-

In Eq. (5), *m*_*G*_ is the efficacy of the GLUT1 inhibitor, and *m*_*A*_ is the efficacy of VEGF

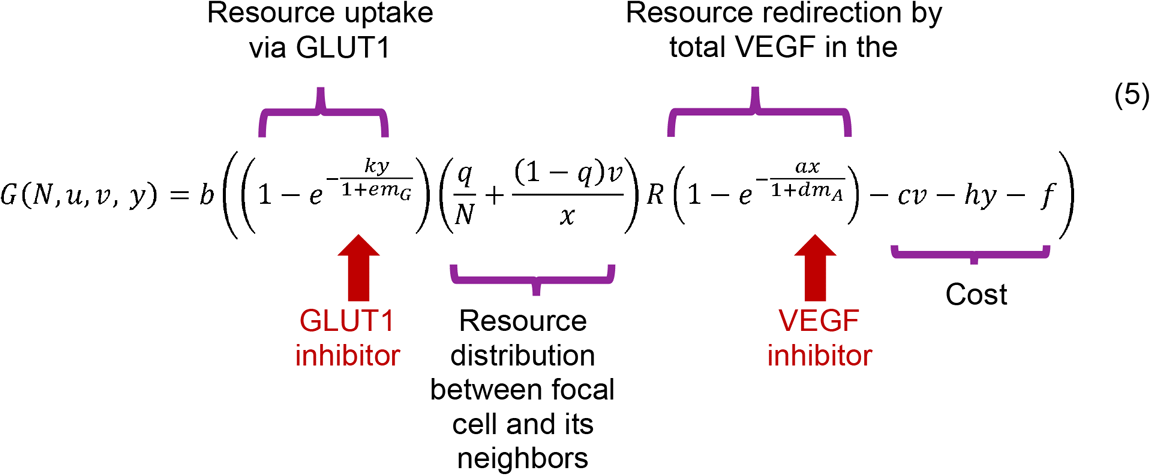

inhibitor.

### 2.2 Simulations

All simulations were performed in Python. To identify the evolutionarily stable strategy (ESS) and the team optimum, the system was iterated until equilibrium was reached, and the resulting steady-state values were recorded. Details of the sensitivity analysis are provided in the Supplementary Section. Variables and parameters used in the model are summarized in Tables 1 and 2.

**Table 1:** Summary of variables and their symbols used in the model.

| Symbol | Meaning |
| --- | --- |
| $v$ | VEGF produced by focal cell |
| $u$ | Average VEGF produced by all cells in a neighborhood |
| $x$ | Total VEGF produced by all cells in a neighborhood |
| $N$ | Number of cells in a neighborhood |
| $y$ | Average GLUT1 expression by cancer population |

**Table 2:**
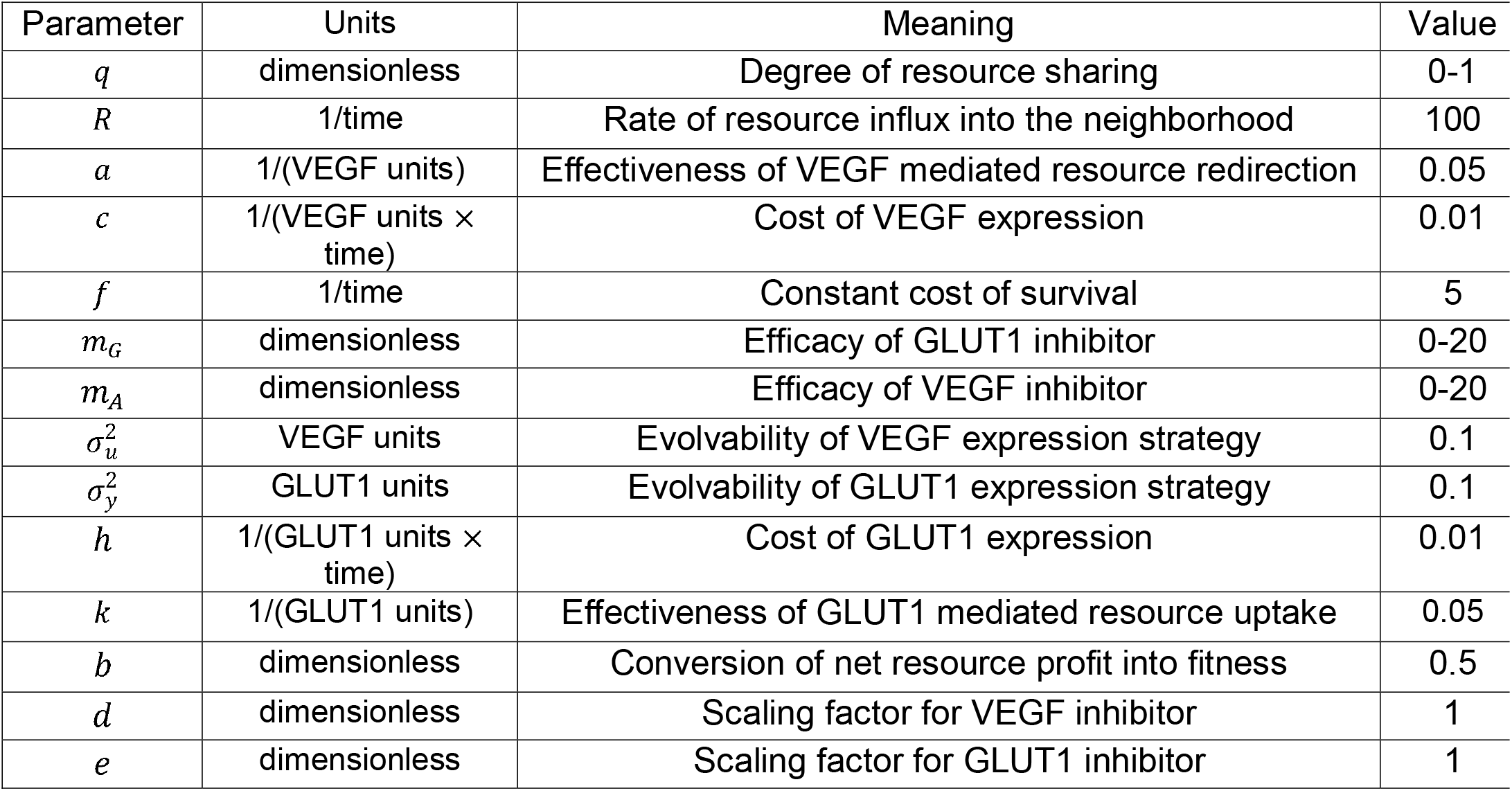
Summary of parameters and parameter values used in the model.

In this theoretical body of work, we are interested in the emergent model dynamics in different parameter ranges. The scaling and units of the parameters can be adjusted to align with experimental measurements. For example, VEGF may be quantified using tissue-level RNA expression or circulating protein concentration, while GLUT1 expression may be measured via RNA expression or immunofluorescence intensity [72,74–77].

## 3. Results

We use an evolutionary game theoretic framework [59,64] to examine how cancer cells allocate investment between VEGF and GLUT1 expression. We examine and compare the team optimum and the evolutionarily stable strategy (ESS) [21,55]. The team optimum is the strategy that maximizes the collective fitness, whereas the evolutionary stable strategy (ESS) is the strategy that, when common among the cancer cells cannot be invaded by rare alternative strategies. In our model, the ESS will be expected outcome of natural selection.

### 3.1 Fixed Neighborhood

We begin by assuming that the number of cancer cells within a neighborhood (*N*) is fixed, and analyze how VEGF and GLUT1 expression evolve across neighborhoods of different sizes.

#### 3.1.1 Team Optimum

For the team optimum, we determine the collective value of VEGF (*u*) and the individual value of GLUT1 (*y*) that maximizes fitness, *G*, for a certain neighborhood size (*N*). We start by setting *v* = *u* in Eq. (1):

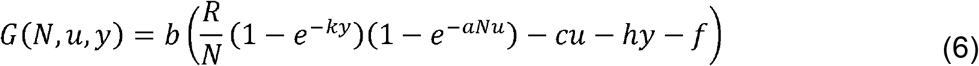

At the fitness-maximizing strategies, the partial derivatives of *G* with respect to each trait *u* and *y* must equal zero:

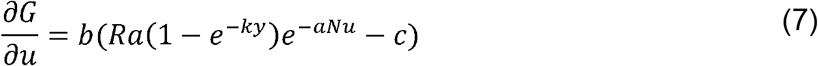

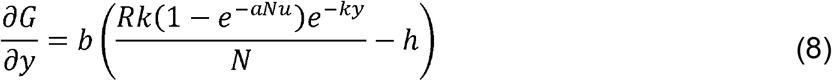

Yielding the following necessary conditions, respectively:

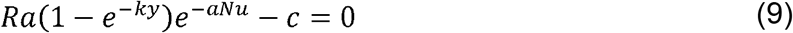

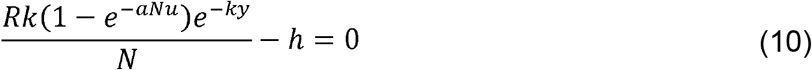

The overall team optimum (*u*_*team*,_ *y*_*team*_) is obtained *numerically* by solving Eq. (9) and (10).

As we have noted before [21], the team optimum is independent of the resource-sharing parameter *q*. Also, the team optimum is independent of the rate at which net profit can be converted into survivorship and proliferation, *b*, and independent of the cell’s fixed cost, *f*. We see how the two traits become co-adapted in that increasing GLUT1 expression increases the marginal value of VEGF production and vice-versa.

Fig. S1 shows how the team-optimum varies with model parameters in fixed neighborhoods. Both per cell VEGF and GLUT1 expression (*u*_*team*_ and *y*_*team*_ respectively) increase with greater resource availability, *R* and decrease with increasing neighborhood size, *N*, and higher expression costs, *c* and *h*. Interestingly, increasing the effectiveness of VEGF-mediated resource redirection, *a*, decreases the team-optimum VEGF expression while increasing GLUT1 expression, whereas the opposite pattern is observed when GLUT1-mediated uptake efficiency, *k*, increases. These trends arise because greater functional effectiveness reduces the level of investment required in that strategy to achieve the same benefit. Fig. S1B shows that VEGF expression at the team optimum is most strongly influenced by neighborhood size, *N*, followed by VEGF effectiveness, *a*, whereas GLUT1 expression is most sensitive to resource availability, *R*, neighborhood size, *N*, and GLUT1 efficiency, *k*.

Fig. 1 shows the temporal dynamics of per cell VEGF (*u*) and GLUT1 (*y*) expression as the system adapts towards the team optimum for five different neighborhood sizes, increasing from left to right. Since we are interested in relative changes in VEGF and GLUT1 expression in response to parameter variation, we chose parameters such that, at *N* = 1, VEGF and GLUT1 expressions are equal. Accordingly, we set the costs and efficacies of VEGF and GLUT1 to be the same, i.e., *c* = *h* and *a* = *k*. Thus, in the absence of neighbors, the focal cell invests equally in VEGF and GLUT1. As neighborhood size increases, investment in both traits declines. This occurs because, as the number of cells in the neighborhood increases, VEGF concentration increases, reducing the marginal benefit of further investment in VEGF. At the same time, the amount of resources available per cell decreases, lowering the payoff from investing in GLUT1. Note that across all neighborhood sizes, the team optimum consistently favors higher investment in GLUT1 relative to VEGF. Although individual investment in VEGF and GLUT1 decreases with increasing neighborhood size, total VEGF expression always increases with neighborhood size (Fig. S2).

**Fig. 1:**
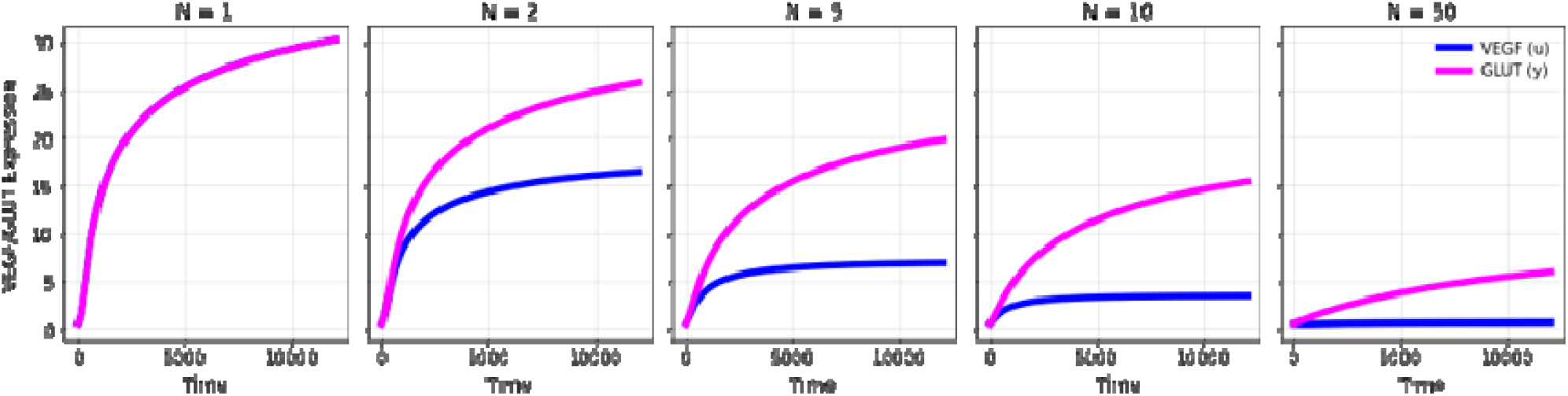
Effect of neighborhood size (*N*) on VEGF and GLUT1 expression at team optimum.

#### 3.1.2 Evolutionary Stable Strategy

The ESS VEGF expression, *u*_*ESS*_, occurs when *v = u*_*ESS*_ maximizes an individual’s fitness given the VEGF production of others. At the ESS, when all cells adopt the same strategy, no alternative strategy can invade. The first order necessary condition for ESS values of *u* and *v* occur when the selection gradients in *v* and *y* equal zero when evaluated at *v = u*_*ESS*_ [78]:

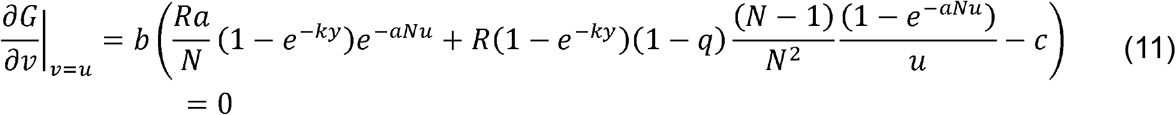

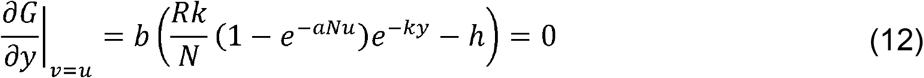

For GLUT1, the ESS condition, Eq. (12), is identical to the team optimum condition, Eq. (10), and both are independent of *q*. This is because GLUT1 expression is purely density dependent and does not depend on the strategies of neighboring cells. For VEGF, Eq. (11) has more terms than the condition for the team optimum, Eq. (9). But when there is complete equal sharing (*q =* 1) or when the neighborhood size is just one (*N* = 1) then the conditions for the ESS become the same as the team optimum. Thus when *q* = 1 or *N* = 1: *u*_*ESS*_ = *u*_*team*_ and *y*_*ESS*_= *y*_*team*_.

Consistent with this, Fig. S3A shows that increasing the degree of resource sharing leads to a decrease in VEGF expression at the ESS, while GLUT1 expression remains relatively unchanged. This pattern is also evident in Fig. 2 and S4: for a fixed neighborhood size, varying *q* has an imperceptible effect on GLUT1 expression, whereas increased resource sharing consistently reduces VEGF expression at the ESS.

**Fig. 2:**
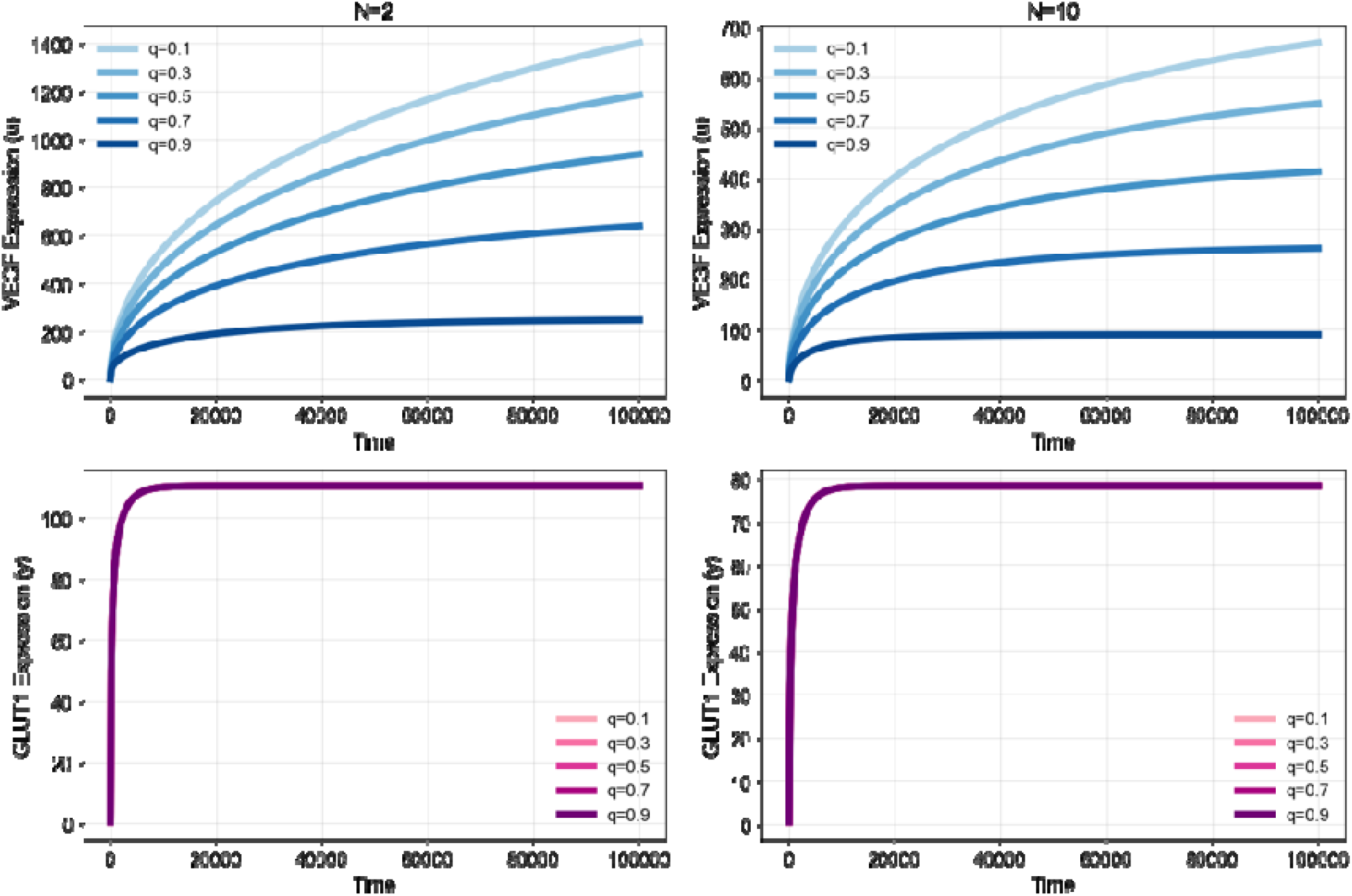
Evolution of VEGF and GLUT1 expression towards the ESS in two neighborhood sizes (*N =* 2 and *N =* 10 ) for different degrees of resource sharing (*q*).

Fig. S5 depicts the adaptive landscape of a focal cell introduced into neighborhoods of varying sizes (1-50). In each landscape, the red star marks the evolutionary stable strategy (ESS), and the black dots trace the trajectory of the population towards that point. When resource sharing is low, equilibrium VEGF expression is substantially higher than when sharing is high. However, in both regimes, increasing neighborhood size consistently reduces the equilibrium values of both VEGF and GLUT1. As neighborhood size (*N*) increases, the entire landscape progressively sinks toward zero fitness and becomes increasingly flat. This indicates that large neighborhoods confer little selective advantage to any single cell. In other words, when many cells share the same resource pool, fitness differences across strategies vanish, as both VEGF and GLUT1 production go to 0.

In Fig. 3 and S3 we examine the effect of neighborhood size on the ESS. We find that with increasing neighborhood size, VEGF expression at the ESS initially increases. This is because for smaller neighborhoods, cells have more to gain by co-opting resources from their neighbors. This incentive vanishes as the neighborhood size increases. Once there are too many cells in the neighborhood, there is little fitness to gain by producing more VEGF. Thus, after a certain point, VEGF expression decreases with increasing neighborhood size. GLUT1 expression decreases smoothly with increasing neighborhood size (in all three resource-sharing environments, see Fig. S3B, C, D), reflecting the actual harvestable resources available to a cell. As before, total VEGF and GLUT1 expressions increase with neighborhood size (Fig. S6).

**Fig. 3:**
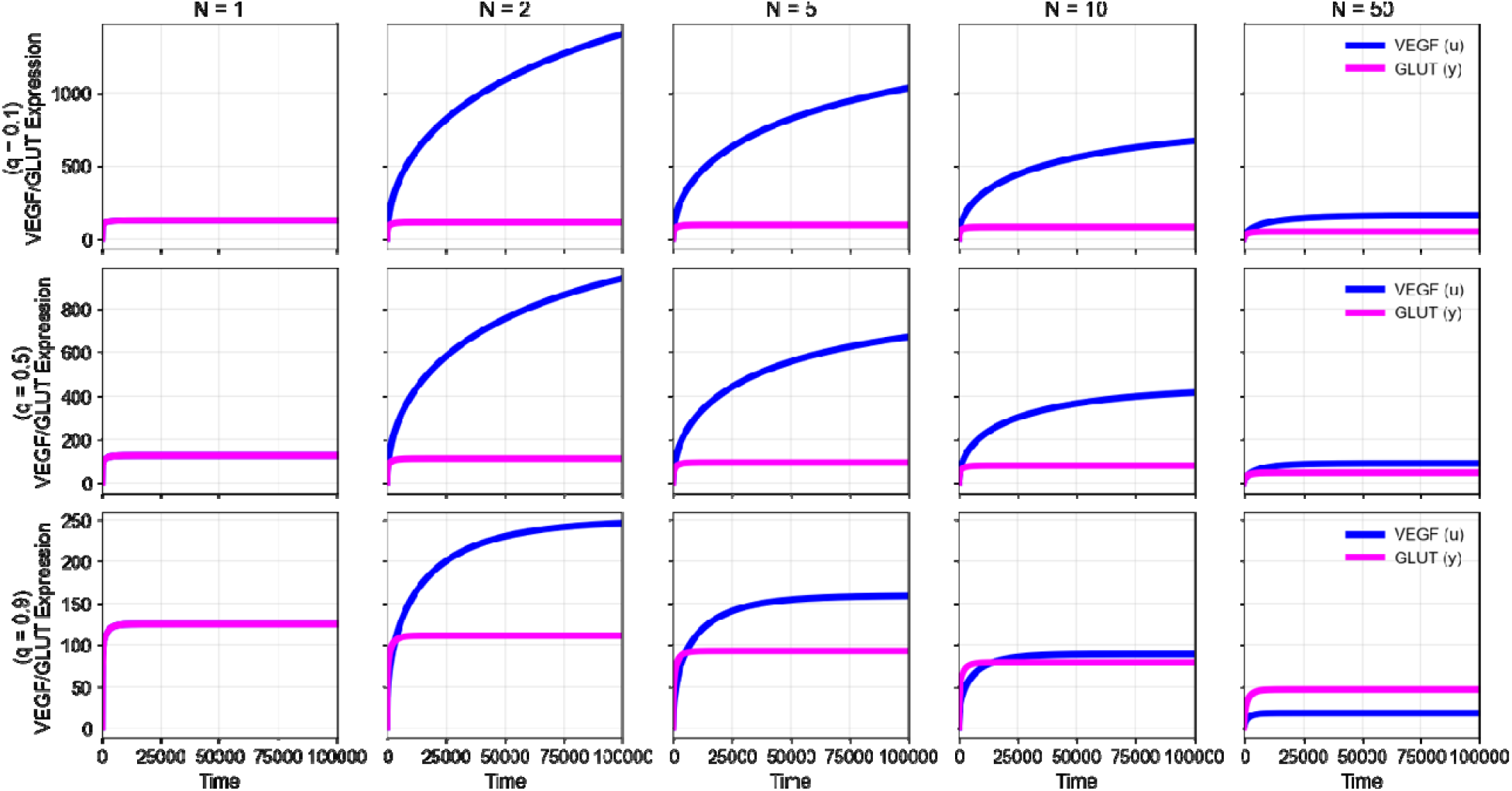
Dynamics of a cell’s VEGF and GLUT1 expression towards the ESS for high (*q* = 0.9), intermediate (*q* = 0.5), and low (*q =* 0.1) resource sharing in neighborhoods of different sizes (*N =* 1 to 50)

As neighborhood size increases from 1 to 2 cells, VEGF expression increases across all three resource-sharing conditions (Fig. 3). This reflects the introduction of direct competition, which leads to an “arms race” as cells invest more in VEGF to secure resources [79]. With further increases in neighborhood size, VEGF expression declines, as the benefits of additional investment in VEGF are shared among neighbors. The neighborhood size at which per-cell VEGF expression (*u*_*ESS*_ ) begins to decline with *N* occurs at lower *N* under high resource sharing. Regardless *u*_*ESS*_ is substantially higher under low resource sharing than under high resource sharing.

GLUT1 expression also decreases with increasing neighborhood size, although this decline is more gradual than for VEGF. This reduction becomes more pronounced under higher resource sharing. Overall, VEGF expression is greater in smaller neighborhoods with low resource sharing, whereas GLUT1 expression dominates in larger neighborhoods with high resource sharing. The interplay between neighborhood size and resource sharing is illustrated in Fig. 3. In large neighborhoods (*N* = 50), VEGF expression is more dominant under low resource sharing (*q* = 0.1), whereas under high resource sharing (*q* = 0.9), GLUT1 expression becomes the strategy in which cells invest more heavily. In contrast, in very small neighborhoods (*N* = 2), VEGF expression remains higher even under high resource sharing conditions, reflecting the strong incentive for cells to invest in VEGF in small neighborhoods. These patterns reinforce that VEGF and GLUT1 are not independent strategies, but coadapted responses to local cellular interactions and resource distribution.

To analyze how these dynamics change with model parameters, we performed parameter sweeps (Fig. S3) and sensitivity analyses (Fig. S7). Both VEGF and GLUT1 expression increase with resource abundance, *R*. Over the range of values considered, variation in VEGF efficacy, *a* has minimal effects on the ESS, whereas GLUT1 efficacy, *k*, strongly influences system behavior. Increasing *k* leads to reduced GLUT1 expression at the ESS, accompanied by increased VEGF expression. Increasing the costs of VEGF, *c*, or GLUT1, *h*, reduces expression of both traits; these reductions are more pronounced under high resource-sharing conditions.

Sensitivity analysis further reveals that GLUT1 expression at the ESS is most sensitive to changes in the efficacy of transporters, *k*, whereas VEGF expression is most sensitive to neighborhood size, *N*, resource abundance, *R*, and the cost of VEGF, *c*, particularly under low resource sharing. In high resource-sharing conditions, VEGF expression is most strongly influenced by the degree of sharing, *q*.

### 3.2 Integrating Ecological and Evolutionary Dynamics

In the previous section, the number of cells in a neighborhood (*N*) was fixed. Here, we let the number of cells in the number increase or decrease based on whether fitness is positive (*G* > 0) or negative (*G* < 0). The ESS will now include VEGF and GLUT1 expression and an equilibrium neighborhood population size, *N*_*ESS*_. The value *u*_*ESS*,_ *y*_*ESS*_ and *N*_*ESS*_ now represent eco-evolutionary dynamics.

We can visualize the eco-evolutionary dynamics on adaptive landscapes (Fig. S8). Each graph is a snapshot in time in the trajectory of the population towards eco-evolutionary stability. The condition for eco-evolutionary stability is attained when Eq. (2), (3), (4) are all satisfied. The black dots show the trajectory and the red star is the ESS strategy itself. As in the fixed-neighborhood case, higher resource sharing leads to lower investment in both VEGF and GLUT1 at the ESS, and to a higher equilibrium population size.

Fig. 4A shows that population size, *N*_*ESS*_ increases monotonically with the degree of resource sharing (*q*), while VEGF and GLUT1 expression, *u*_*ESS*_ and *y*_*ESS*_, are inversely related to *q* (also see Fig. S9). Higher resource sharing reduces the escalating VEGF-expression arms race that occurs at low resource sharing as cancer cells attempt to secure a higher share of the bloodborne resources. Figure 4B further illustrates how determines the relative balance between VEGF and GLUT1 expression within a neighborhood. Under low resource-sharing conditions, the ratio of VEGF to GLUT1 expression is high, reflecting the strong incentive for individual cells to invest in VEGF-mediated signaling. In contrast, high resource-sharing environments reduce this ratio, favoring relatively greater investment in GLUT1 expression (also see Fig. S10).

**Fig. 4:**
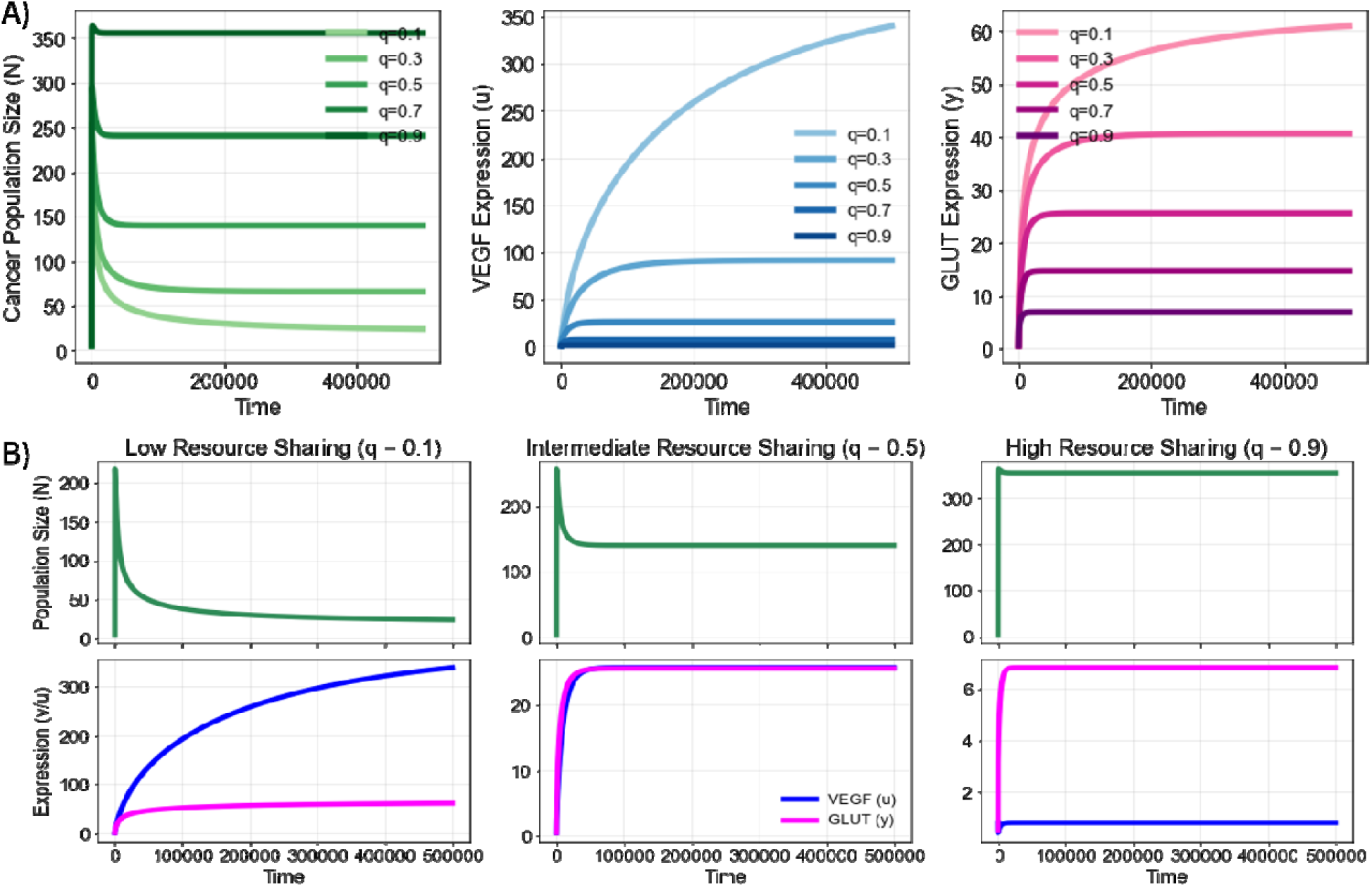
A) Population Size (*N*), VEGF Expression (*u*), and GLUT1 Expression (*y*) dynamics for different values of *q*. Darker the shade, higher the resource sharing (*q*). B) Comparison of ESS dynamics in low (*q* = 0.1), intermediate (*q* = 0.5), and high (*q* = 0.9) resource sharing neighborhoods.

To characterize how ecological and evolutionary interactions shape neighborhood dynamics, we examine how key model parameters influence both the ESS and team optimum through parameter sweeps and sensitivity analyses (Fig. S11-S14). Across all regimes, increasing resource abundance, *R*, increases population size. At the ESS, VEGF and GLUT1 expression increase with *R* under low and intermediate resource sharing, whereas under high resource sharing, both traits exhibit non-monotonic behavior, initially decreasing before increasing at higher resource levels. In contrast, at the team optimum, increasing *R* leads to monotonic reductions in VEGF and GLUT1 expression, reflecting reduced need for costly resource acquisition.

Increasing the efficacy of GLUT1-mediated uptake, *k*, increases population size while reducing both VEGF and GLUT1 expression at the ESS, reflecting more efficient resource acquisition and utilization. This effect is consistent across low, intermediate, and high resource-sharing conditions, as well as at the team optimum. In contrast, variation in VEGF efficacy, *a*, has little effect on the ESS across all resource-sharing regimes. But at the team optimum, increasing *a* increases population size while reducing investment in both VEGF and GLUT1 expression.

The effect of increasing the cost of VEGF, *c*, depends on the degree of resource sharing. Under low resource sharing, VEGF expression decreases sharply with increasing *c*, with little effect on the population size or GLUT1 expression. At intermediate resource sharing, both VEGF and GLUT1 expression decline with increasing *c*, while population size remains nearly constant. Under high resource sharing, increasing *c* reduces both population size and VEGF expression, with only little change in GLUT1. At the team optimum, increasing *c* reduces population size and VEGF expression while increasing GLUT1 expression, consistent with compensatory allocation toward GLUT1.

Increasing the cost of GLUT1 expression, *h*, has a consistent effect across all resource-sharing regimes: population size and GLUT1 expression decline, while VEGF expression increases, indicating a compensatory shift away from costly GLUT1 investment and toward VEGF. The same qualitative pattern is also observed at the team optimum. Finally, increasing the cost of survival, *f*, reduces population size while increasing investment in both VEGF and GLUT1 across all resource sharing values and at the team optimum.

Normalized sensitivity analysis (Fig. S15) identifies the dominant drivers of system behavior. At the team optimum, population size is most strongly influenced by resource abundance, *R*, and GLUT1 efficacy, *k*. VEGF expression is primarily governed by the cost of GLUT1 expression, *h*. At the ESS, the sensitivities depend on the degree of resource sharing. Under low and intermediate sharing, VEGF expression is most sensitive to VEGF cost, *c*, whereas GLUT1 expression is primarily driven by GLUT1 efficacy, *k*. In high resource sharing, VEGF expression becomes overwhelmingly sensitive to the degree of sharing, *q*, and other parameters become less impactful.

### 3.3 The Effect of Therapy

In this section, we examine the effects of two potential therapeutic strategies: one targeting VEGF signaling and one targeting GLUT1-mediated glucose uptake[67,72,80]. The VEGF-targeting therapy, *m*_*A*_, acts by reducing the efficacy of VEGF-mediated resource redirection, *a*, while the GLUT1-targeting therapy, *m*_*G*_, reduces the efficacy of glucose uptake, *k*. We expect either therapy to induce predictable coadapted phenotypic responses in both VEGF and GLUT1 expression. As in Sections 3.1 and 3.2, we analyze both the team optimum and evolutionarily stable strategy (ESS) under these interventions. Details of the derivations are provided in the Supplementary material.

Considering the both therapies, *m*_*A*_ and *m*_*G*_, the necessary conditions for the team optimum VEGF and GLUT1 expression, *u*_*team*_ and *y*_*team*_, respectively are

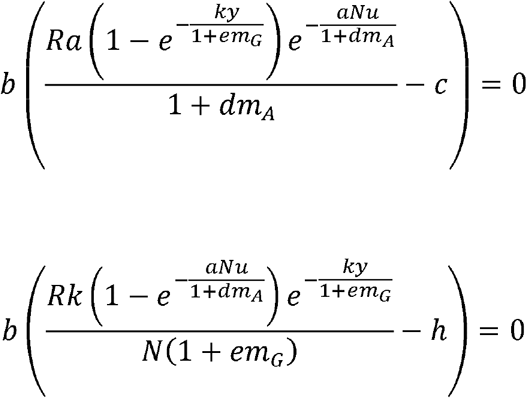

Under therapy, the ESS conditions for VEGF and GLUT1, *u*_*ESS*_ and *y*_*ESS*_ are:

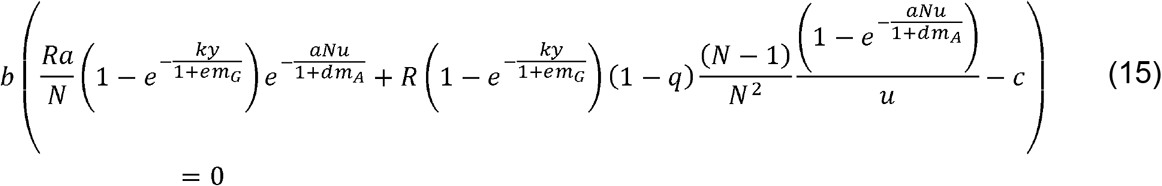

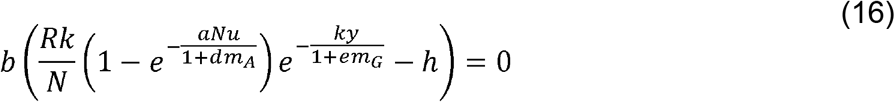

For either the team optimum or the ESS, the condition for the neighborhood’s equilibrium population size, *N*_*team*_ and *N*_*ESS*_, respectively, is:

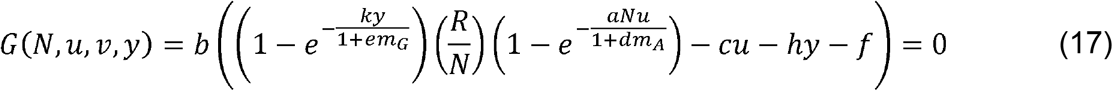

Eco-evolutionary stability is attained when Eq. (15), (16), and (17) are satisfied.

As shown in Fig. 5, inhibition of GLUT1 consistently reduces tumor size more effectively than anti-angiogenic therapy across all degrees of resource sharing. In our framework, this happen because GLUT1-mediated uptake operates at cellular level and is largely independent of the degree of resource sharing (*q*). In contrast, the effectiveness of anti-angiogenic therapy is heavily influenced by *q*. VEGF inhibition is most effective under high resource-sharing conditions (*q* = 0.9) and has minimal impact under low resource-sharing regimes. While one might expect VEGF inhibitors to be more effective in tumors with higher VEGF expression, our results suggest that the tragedy of the commons that emerges under low resource sharing buffers the impact of anti-angiogenic therapy. This could mean that therapies targeting shared paracrine factors may be less effective than those targeting cell-autonomous mechanisms such as glucose uptake.

**Fig. 5:**
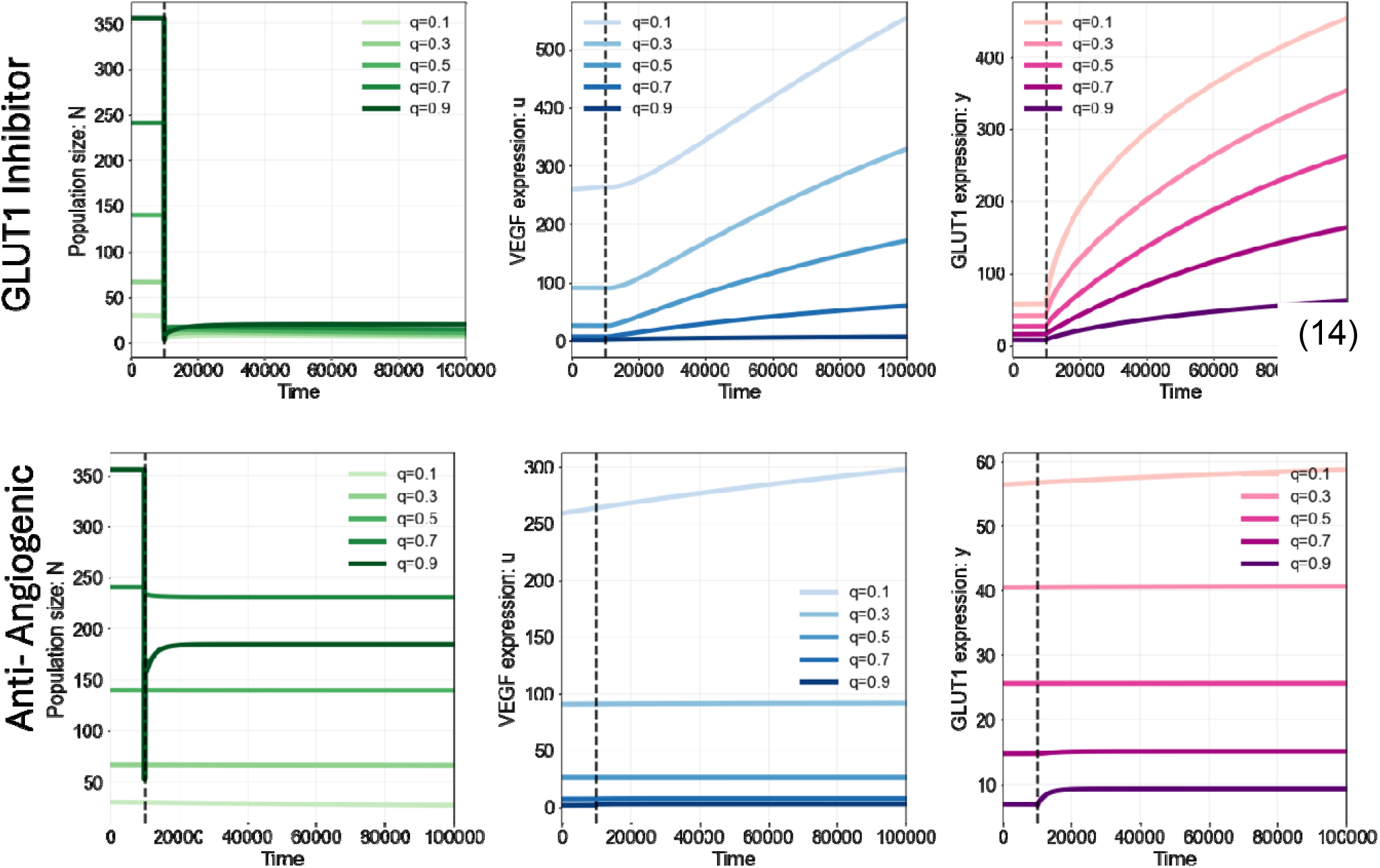
Effect of GLUT1 inhibitor (top) and anti-angiogenic therapy (bottom) on population size (*N*), VEGF expression (*u*), GLUT1 expression (*y*) for different degrees of resource sharing (*q*). For the top row: *m*_*G*_ = 10 and *m*_*A*_ = 0. For the bottom row, *m*_*G*_ = 0 and *m*_*A*_ = 10. Therapy is administered at *t* = 10000 (dashed line).

We further examine these differences in Fig. 6, which shows the effect of varying therapy doses across low, intermediate, and high resource-sharing environments. As expected, increasing the dose generally reduces tumor size. However, anti-angiogenic therapy remains largely ineffective in low and intermediate resource-sharing regimes, whereas both therapies are more effective under high resource sharing. These results underscore that therapeutic response is shaped not only by drug potency, but also by the interactions between cells in a tumor.

**Fig. 6:**
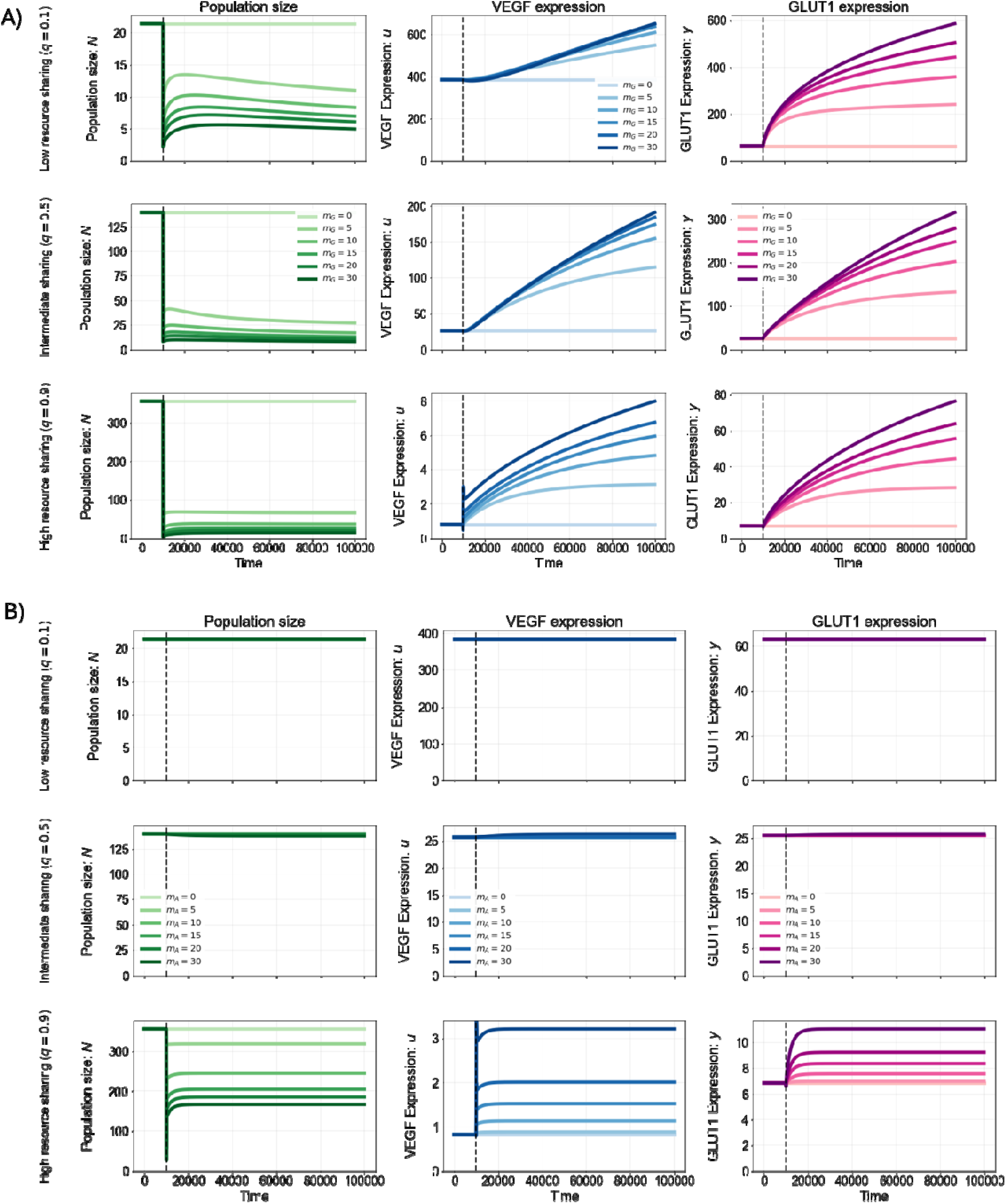
Effect of different doses of (A) GLUT1 inhibitor (*m*_*G*_) and VEGF inhibitor (*m*_*A*_) (B) on population size (*N*), VEGF expression (*u*), GLUT1 expression (*y*) at ESS. For the top row: *q* = 0.1 (low resource sharing), middle row: *q* = 0.5 (intermediate resource sharing), bottom row: *q* = 0.9 (high resource sharing). Therapy is administered at *t* = 10000 (dashed line).

Fig. S16 summarizes the percentage decrease in population size following monotherapy across the three resource-sharing regimes. Fig. 7 examines the combined effect of VEGF and GLUT1 inhibition. Relatively low doses of GLUT1 inhibition (*m*_*G*_ *=* 5) achieve reductions in tumor size that are not attainable even with high doses of VEGF inhibition (*m*_*A*_ = 50) alone. Moreover, combining moderate anti-angiogenic therapy with lower doses of GLUT1 inhibition yields comparable reductions in tumor size. These results suggest that combination strategies can achieve substantial therapeutic benefit at reduced dosing, which may be particularly important given the systemic constraints associated with targeting glucose uptake.

**Fig. 7:**
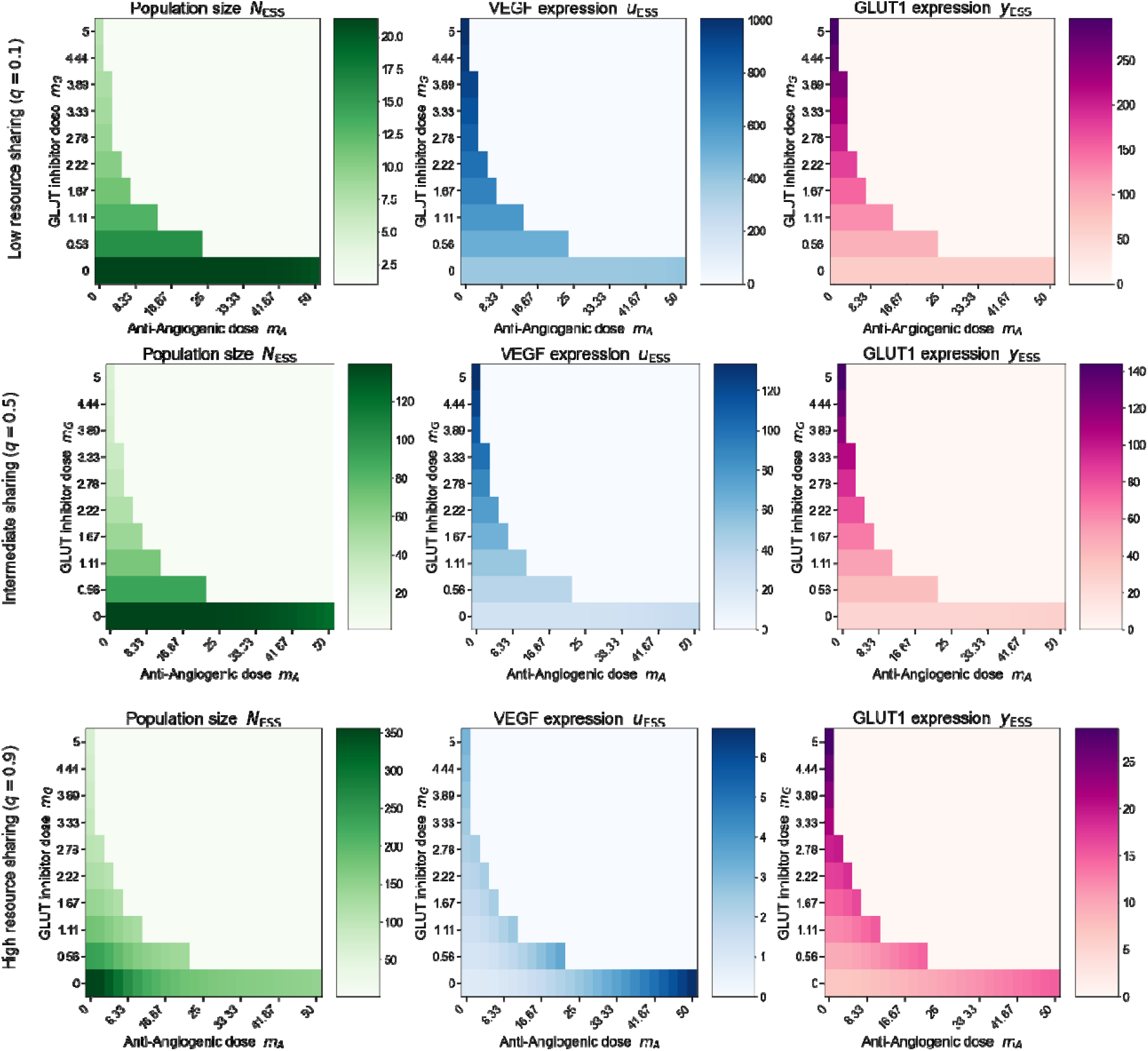
Effect of combination therapy with GLUT1 inhibitor (*m*_*G*_) and VEGF inhibitor (*m*_*A*_) on population size (*N*), VEGF expression (*u*), GLUT1 expression (*y*) at ESS. For the top row: *q* = 0.1 (low resource sharing), middle row: *q* = 0.5 (intermediate resource sharing), bottom row: *q* = 0.9 (high resource sharing).

Our analysis suggests that therapeutic response is heavily influenced by the degree of resource sharing within the tumor microenvironment. The degree of resource sharing is an empirical question. This parameter may vary by disease type, patient to patient, or even among different regions of the same tumor, leading to spatial variation in the evolutionary incentives governing VEGF and GLUT1 co-expression. Consequently, the effectiveness of a given therapy may differ across tumor neighborhoods, even within a single lesion. By framing VEGF and GLUT1 expression as coadaptations shaped by neighborhood structure, our model provides a foundation for designing therapies that anticipate evolutionary responses rather than react to them [81,82].

## 4. Discussion

In this study, we examined the coadaptation [83,84] of VEGF-mediated angiogenesis and GLUT1-mediated glucose uptake as complementary foraging strategies in cancer cells [56]. Both traits are downstream effectors of HIF-1, which is activated in response to resource deprivation and hypoxia, linking the two coadaptations with a shared regulatory program [9–11,24,26]. Clinical and translational studies have consistently documented coordinated expression of hypoxia-related markers, including VEGF, GLUT1, CA-IX, and HIF-1α, across multiple malignancies [29,31]. In cancer, both VEGF and GLUT1 are frequently overexpressed relative to normal tissue. Immunohistochemical analyses in cervical, ovarian, gastric, and bladder cancers demonstrate frequent co-expression and spatial co-localization of these markers, often correlating with tumor progression, microvessel density, and poor prognosis [25,27,28,85–87]. Although these studies do not explicitly address evolutionary dynamics, they provide strong biological evidence that angiogenic and metabolic responses are activated together in human tumors.

A substantial mathematical oncology literature has modeled angiogenesis and tumor metabolism, but typically in isolation. Mechanistic models of angiogenesis have focused on VEGF dynamics, endothelial sprouting, vascular remodeling, and nutrient transport using reaction–diffusion, continuum, and multiscale frameworks to predict tumor growth and vascular architecture [34,35,88]. Separately, models of tumor metabolism have examined glucose uptake, glycolytic flux, and metabolic competition, and recent evolutionary game-theoretic work has treated GLUT1 expression as a tragedy of the commons [41,44–46,55]. Our framework explicitly models VEGF and GLUT1 as coadapted traits whose joint evolutionary dynamics are shaped by neighborhood interactions via resource supply and degree of resource sharing.

VEGF influences resource availability at the neighborhood scale, whereas GLUT1 governs resource uptake at the cellular scale. Modeling these traits jointly reveals that their evolutionary dynamics are coupled and shaped by cell–cell interactions, specifically the degree of resource sharing. Here, resource sharing refers to how resources supplied by blood vessels are distributed among cells within a local habitat. We show that the degree of resource sharing within tumor neighborhoods is a key determinant of strategy selection [21]. Under low-resource sharing, cells are incentivized to overinvest in VEGF, leading to a tragedy of the commons that reduces collective tumor fitness [54,55]. This mirrors ecological systems in which shared resources are overexploited when access is unequal [89–93]. In contrast, when resources are more equitably distributed, the selective pressure to overproduce VEGF diminishes, and cellular investment shifts toward uptake efficiency through GLUT1 expression. Thus, resource sharing dynamics influences the dominant foraging strategy in a neighborhood.

GLUT1 reflects a cell’s capacity for resource uptake, whereas VEGF reflects the ‘effort’ per resource recruited into the neighborhood. The ratio of VEGF to GLUT1 expression, thus, provides a proxy for how much effort is expended on resource recruitment relative to uptake. A high VEGF-to-GLUT1 ratio suggests environments where cells must invest heavily in recruiting resources that are not readily shared, consistent with low resource-sharing conditions. Conversely, a lower ratio indicates that resources are more accessible, allowing cells to prioritize uptake over blood vessel recruitment.

Different tumor regions are likely to exhibit distinct degrees of resource sharing depending on the resource supply, vascular density, hypoxia, and cell packing [94–96]. Consequently, some neighborhoods may favor VEGF and others GLUT1 expression. To empirically estimate resource-sharing dynamics in cancer, multiplex imaging platforms such as immunofluorescence and spatial transcriptomics can be used to simultaneously quantify multiple tumor markers at single-cell resolution while preserving spatial context [97,98]. This spatial information enables the identification of local neighborhoods, within which marker intensities can be quantified for each cell. From these measurements, the VEGF-to-GLUT1 ratio can be computed at the neighborhood level and used as a proxy for resource-sharing dynamics. Model predictions can be tested by examining how this ratio varies with features such as vascular density and local cell packing. This spatial heterogeneity suggests that therapeutic responses may vary not only between tumors in different patients but also across regions of a single lesion.

Our simulations suggest that targeting glucose uptake exerts a stronger suppressive effect on tumor size than anti-angiogenic therapy across different degrees of resource sharing. Clinically, anti-angiogenic agents such as bevacizumab and other VEGF pathway inhibitors are widely used, yet their benefits are often transient, with tumors frequently developing resistance [99–102]. Several studies have reported that resistance to VEGF-targeted therapies is accompanied by enhanced GLUT1 expression and metabolic reprogramming toward glycolysis [94,103], suggesting a compensatory shift from vascular recruitment to cellular resource acquisition.

In contrast, systemic GLUT1 inhibition is currently not a viable strategy due to the essential role of glucose transport in normal tissues [6,104,105]. Several GLUT inhibitors have been explored in preclinical stages, including small molecules such as WZB117, BAY-876, and STF-31 [70,71,73], which inhibit GLUT1-mediated glucose transport and have demonstrated anti-tumor effects in cell lines and xenograft models. Metabolic analogues such as 2-deoxy-D-glucose (2-DG) have also been explored as glycolytic inhibitors [106], though systemic toxicity and limited efficacy have constrained clinical translation. The widespread expression of GLUT1 in erythrocytes and the blood– brain barrier raises significant concerns about on-target toxicity, limiting the therapeutic window of direct GLUT1 blockade [6,107].

Emerging precision-delivery approaches offer potential strategies to overcome these limitations by selectively targeting tumor metabolism. Recent preclinical studies have demonstrated tumor-biased delivery of GLUT1-targeting payloads, including Glut1 siRNA delivered via engineered nanoparticles [108,109], as well as nanoformulations carrying GLUT1 inhibitors such as BAY-876 [110,111]. More broadly, advances in nanomedicine and targeted delivery systems have enabled selective modulation of tumor metabolic pathways while sparing normal tissues [112]. If such tumor-specific targeting can be achieved, our model predicts that GLUT1 inhibition would be highly effective across a wide range of resource-sharing conditions.

An alternative therapeutic approach is to reduce systemic glucose availability rather than inhibiting glucose transport. Dietary interventions such as ketogenic diets, which lower circulating glucose and shift metabolism toward ketone utilization, have shown potential in preclinical models and early clinical studies to impair tumor growth and enhance response to therapy, although their efficacy remains context-dependent and requires further validation [113–115]. Our previous modeling work [21] demonstrated that reducing resource supply is more effective at limiting tumor growth than reducing the efficacy of VEGF-mediated resource recruitment. Potentially, combining reduced resource inflow with inhibition of GLUT1-mediated uptake could be an even more effective therapeutic approach.

Several limitations must be addressed. Our framework is not spatially explicit. Although we consider resource distribution between cells through the parameter *q*, we do not explicitly represent spatial diffusion of oxygen or glucose, or vascular topology. In our model, *q* is treated as a fixed parameter; a natural extension would allow resource sharing itself to evolve or vary dynamically over time.

In addition, we assume that both VEGF and GLUT1 expression incur fitness costs. These costs are biologically plausible. VEGF production requires sustained biosynthetic investment and contributes to structurally abnormal vasculature that may reduce perfusion efficiency and increase metabolic stress. Elevated GLUT1 expression is associated with high glycolytic flux, increased metabolic demand, and altered redox balance, imposing energetic constraints on the cell [57,116–119]. While these costs are represented abstractly in our framework, experimental quantification of the metabolic and physiological trade-offs associated with VEGF and GLUT1 expression would further strengthen evolutionary models of tumor resource acquisition.

In summary, our study frames angiogenesis and glucose uptake as coadapted strategies shaped by ecological interactions within the tumor microenvironment. By modeling VEGF and GLUT1 as coadaptations, we demonstrate that the balance between these two foraging strategies depends on resource sharing and neighborhood structure. These results highlight that tumor behavior and therapeutic response cannot be fully understood without considering how adapted traits interact with each other. More broadly, our findings underscore the importance of integrating ecological structure and evolutionary dynamics into cancer modeling [81]. Viewing tumors as systems of interacting, coevolving strategies may provide a foundation for designing therapies that anticipate evolutionary responses rather than inadvertently selecting for them.

## Supporting information

Supplementary Inormation

## 5. Conflicts of Interest

The authors declare no conflicts of interest.

## 6. Funding

We appreciate grant support from NIH/NCI 1R01CA258089 and 1U54 CA274507-01A1, and DoD W81XWH2210680.

## 7. Author Contributions

RB and JB conceptualized the model and study. RB created the simulations and wrote the manuscript. RB, JB, and RG reviewed and edited the manuscript.

## 8. Acknowledgments

The authors thank Anuraag Bukkuri for his valuable discussions and input in the design of the model.

## 9. Declaration of generative AI and AI-assisted technologies in the writing process

The authors used Grammarly and ChatGPT for copyediting the manuscript. After using the tools, the authors reviewed and edited the content as needed and take full responsibility for the content of the published article.

## 10. Data Availability

Codes used to generate simulations in the paper are available at https://github.com/rbhattacharya49/vegf-glut1-coadaptation

## Notes

https://github.com/rbhattacharya49/vegf-glut1-coadaptation

### Competing Interest Statement

The authors have declared no competing interest.

### Summary of Updates

We have updated the manuscript to include additional figures and corrected errors in equation numbering.

