## Supplementary Inormation for "Modeling VEGF and GLUT1 Expression as Coadapted Foraging Strategies in Cancer"

#### Supplementary Information

Ranjini Bhattacharya [1,2,3], Robert A. Gatenby [1,2,4], Joel S. Brown [1,2]

1 Cancer Biology and Evolution Program, H. Lee Moffitt Cancer Center and Research Institute, Tampa, FL, USA

2 Department of Integrated Mathematical Oncology, H. Lee Moffitt Cancer Center and Research Institute, Tampa, FL, USA

3 Department of Cancer Biology, University of South Florida, Tampa, FL, USA

4 Department of Radiology, H. Lee Moffitt Cancer Center, FL, USA

#### 1. Deriving ESS and Team Optimum for Therapy Case

##### 1.1 Team Optimum under therapy

At the team optimum, all cells adopt a strategy that maximizes the collective fitness of the cancer population. To derive the team optimum conditions, we first set $v = u$ into the fitness generating function $G$ that includes therapy (Eq. 5):

$$G(N,u,y) = b\left(\left(1 - e^{-\frac{ky}{1+em_G}}\right)\frac{R}{N}\left(1 - e^{-\frac{aNu}{1+dm_A}}\right) - cu - hy - f\right) \quad \text{S1}$$

The team optimum, $(u_{team}, y_{team})$ corresponds to the strategy that maximizes $G$. The necessary condition for $(u_{team}, y_{team})$ requires solving two conditions obtained by taking the partial derivatives of $G$ with respect to each trait and setting them equal to zero:

$$\frac{\partial G}{\partial u} = b\left(\frac{Ra\left(1 - e^{-\frac{ky}{1+em_G}}\right)e^{-\frac{aNu}{1+dm_A}}}{1 + dm_A} - c\right) = 0 \quad \text{S2}$$

$$\frac{\partial G}{\partial y} = b\left(\frac{Rk\left(1 - e^{-\frac{aNu}{1+dm_A}}\right)e^{-\frac{ky}{1+em_G}}}{N(1 + em_G)} - h\right) \quad \text{S3}$$

##### 1.2 Evolutionary Stable Strategy with therapy

The evolutionary stable strategy (ESS), $(u_{ESS}, y_{ESS})$ is the best an individual focal cell can do given the environment and strategies of all other cells in the neighborhood. If all cells adopt this strategy, no cell can attain higher fitness by unilaterally changing its strategy At $(u_{ESS}, y_{ESS})$ the partial derivatives of $G$ with respect to $v$ and $y$ must be zero when evaluated at $v = u$:

$$\left.\frac{\partial G}{\partial v}\right|_{v=u} = b\left[\frac{Ra}{N}\left(1 - e^{-\frac{ky}{1+em_G}}\right)e^{-\frac{aNu}{1+dm_A}} + R\left(1 - e^{-\frac{ky}{1+em_G}}\right)(1-q)\frac{(N-1)}{N^2}\frac{\left(1 - e^{-\frac{aNu}{1+dm_A}}\right)}{u} - c\right] = 0 \qquad \text{S4}$$

$$\left.\frac{\partial G}{\partial y}\right|_{v=u} = b\left[\frac{Rk\left(1 - e^{-\frac{aNu}{1+dm_A}}\right)e^{-\frac{ky}{1+em_G}}}{N(1+em_G)} - h\right] = 0 \qquad \text{S5}$$

Note that the ESS condition for GLUT1 expression is the same as the team optimum, while not so for the ESS condition for VEGF. Thus, any difference between $y_{team}$ and $y_{ESS}$ is driven by the difference between $u_{team}$ and $u_{ESS}$.

#### 2. Sensitivity Analysis

To evaluate the robustness of our results, we performed local sensitivity analyses [1] of the equilibrium values $(u_{ESS}, y_{ESS}, N_{ESS})$ and $(u_{team}, y_{team}, N_{team})$ with respect to model parameters

$$p \in \{q, k, a, R, c, h, f\}.$$

In the case of a fixed neighborhood, $N$ is also treated as a parameter.

A baseline parameter set $\theta_0 = \{k_0, a_0, R_0, c_0, h_0, f_0,\ q_0\}$ was first used to compute the reference equilibrium values. Local sensitivities were then estimated using central finite differences. For each parameter $p$, symmetric perturbations were introduced:

$$p_{\pm} = p_0 \pm \Delta p, \qquad \Delta p = \max\left(\alpha \mid p_0 \mid, p_{min}\right),$$

where $\alpha$ denotes a fractional perturbation (e.g., 10%) and $p_{min}$ represents a user-defined minimum absolute perturbation applied to $p_0$ . Parameter bounds (such as $q \in [0,1]$) were enforced where necessary. For each perturbed value, the system was integrated again, and the new equilibrium values were obtained. We approximated finite-difference derivatives as:

$$\frac{dX^*}{dp} \approx \frac{X_+^* - X_-^*}{p_+ - p_-},$$

where $X^*$denotes any equilibrium quantity of interest ($u_{ESS}, y_{ESS}, N_{ESS}$) at ESS or team optimum ($u_{team}, y_{team}, N_{team}$).

To allow comparison across parameters with different scales, normalized sensitivities were computed as

$$S_X = \frac{p_0}{\max(| X_0^* |, \varepsilon)} \frac{dX^*}{dp},$$

with $\varepsilon = 10^{-9}$ to prevent division by zero. These dimensionless sensitivities quantify the relative change in equilibrium outcomes induced by small perturbations in model parameters.

### 3. Supplementary Figures

#### 3.1 Fixed Neighborhood

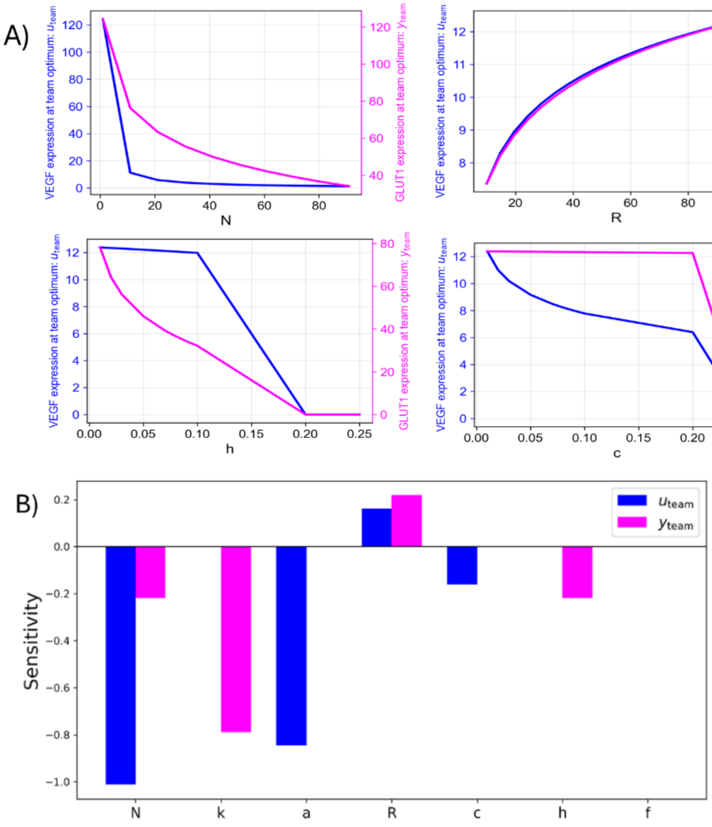


Fig. S1: A) Parameter Sweeps for the team optimum values of VEGF and GLUT1 expression in a fixed neighborhood. The neighborhood size ($N$) is held constant at $N = 10$ for all parameters other than $N$. B) Sensitivity analysis at the team optimum for different parameters.

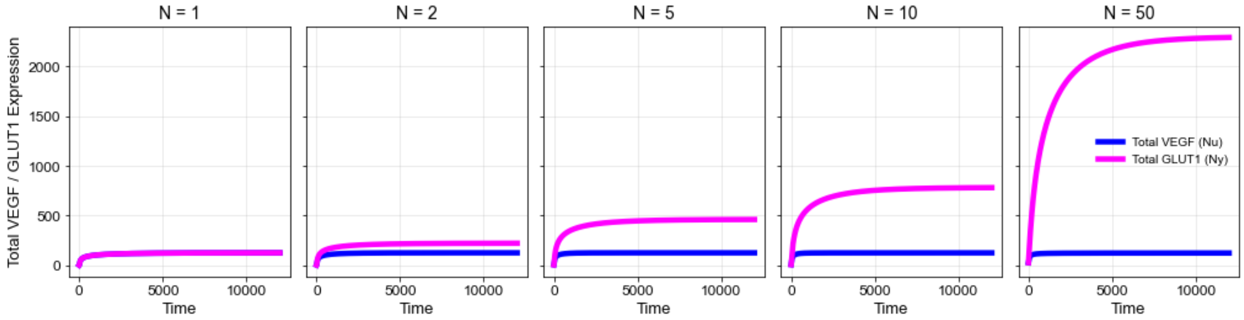


Fig. S2: Effect of neighborhood size ($N$) on total VEGF and total GLUT1 expression at team optimum.

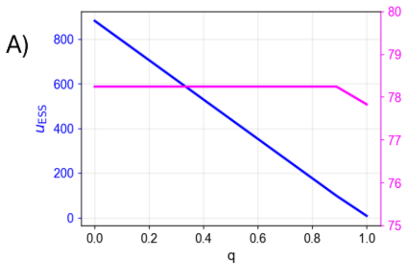


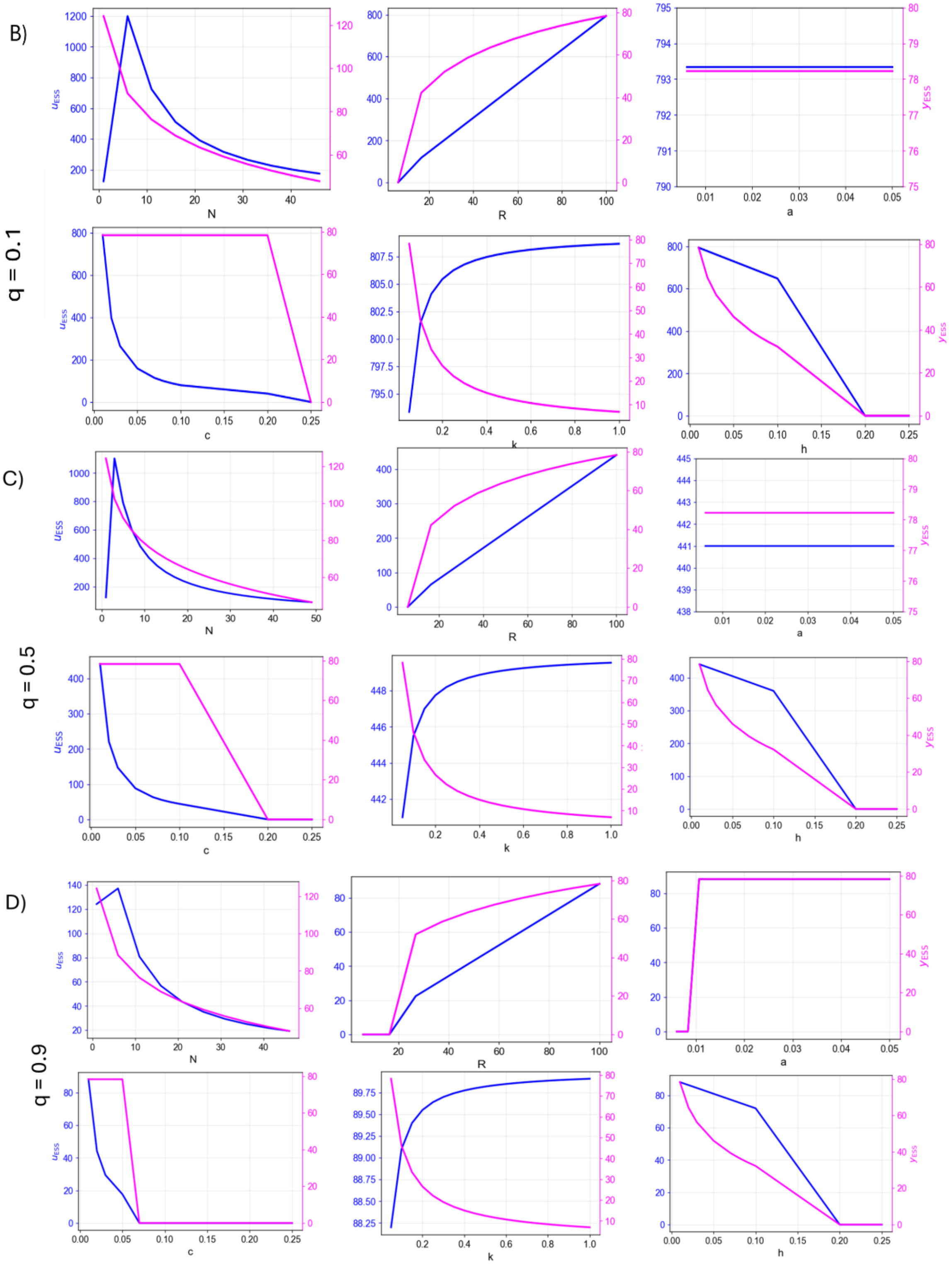


Fig. S3: A) Effect of the degree of resource sharing ($q$) on VEGF and GLUT1 expression at ESS. B), C), D) Parameter Sweeps for VEGF and GLUT1 expression at ESS for low resource sharing ($q = 0.1$), intermediate resource sharing ($q = 0.5$), and high resource sharing ($q = 0.9$), respectively. For all parameter sweeps, the neighborhood size was held at $N = 10$.

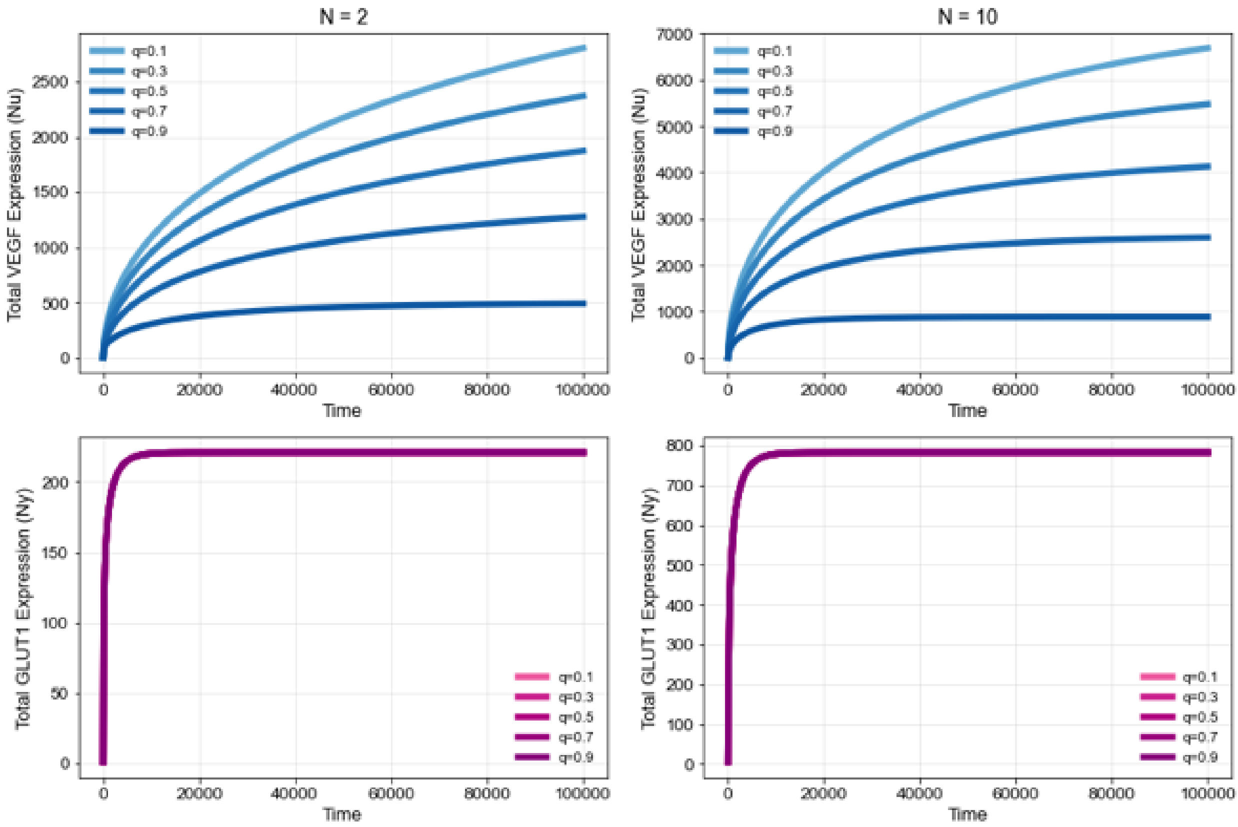


Fig. S4: Evolution of total VEGF and total GLUT1 expression towards the ESS in two neighborhood sizes ($N = 2$ and $N = 10$) for different degrees of resource sharing ($q$).

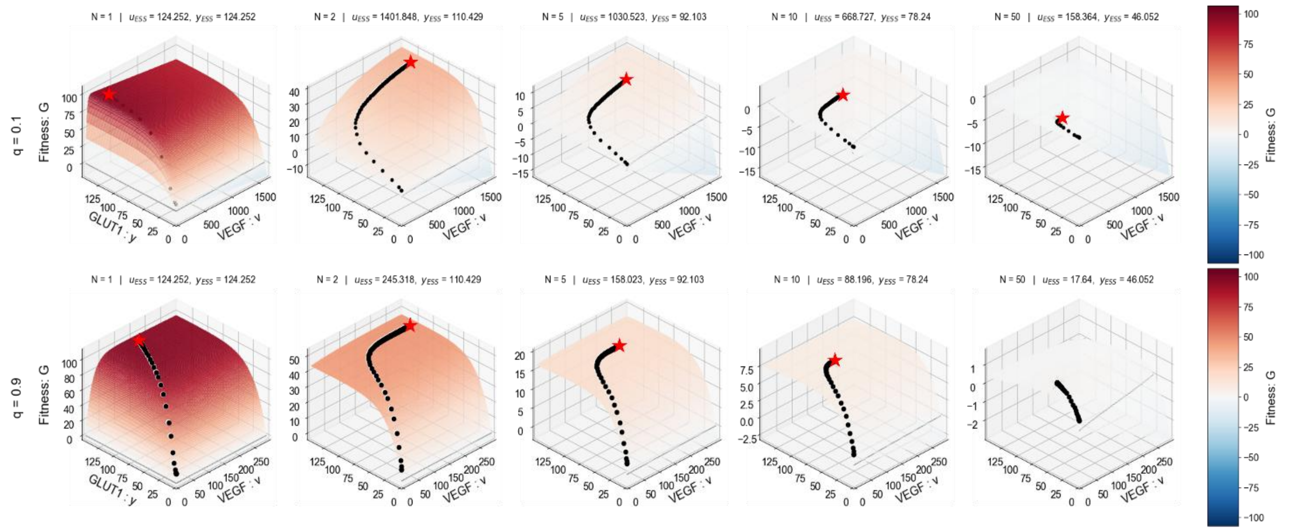


Fig. S5: Adaptive Landscape for Fixed Neighborhood: Each graph represents the adaptive landscape at equilibrium for neighborhoods of different sizes. The red star represents the final trait values at evolutionary equilibrium. The black circles show the path of the population towards the equilibrium. $u_{ESS}$ and $y_{ESS}$ correspond to the equilibrium VEGF and GLUT1 values respectively. Top = Low resource sharing ($q = 0.1$) and bottom = high resource sharing ($q = 0.9$).

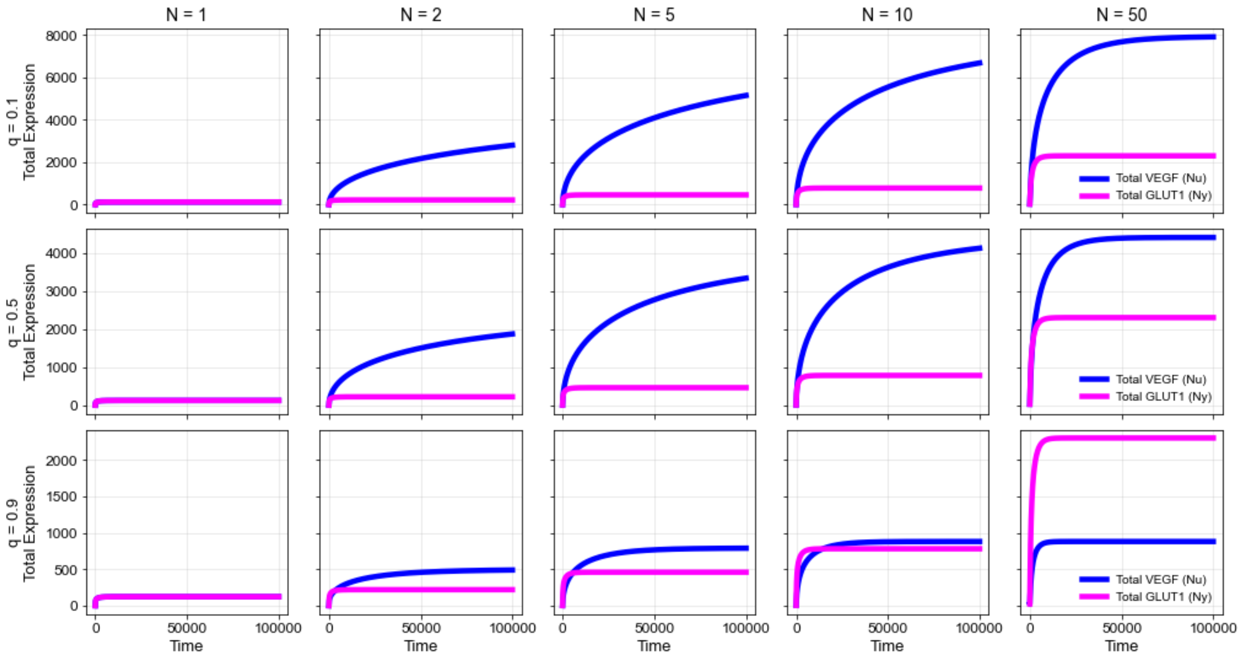


Fig. S6: Dynamics of a total VEGF and total GLUT1 expression at the ESS for high ($q = 0.9$), intermediate ($q = 0.5$), and low ($q = 0.1$) resource sharing in neighborhoods of different sizes ($N = 1$ to 50).

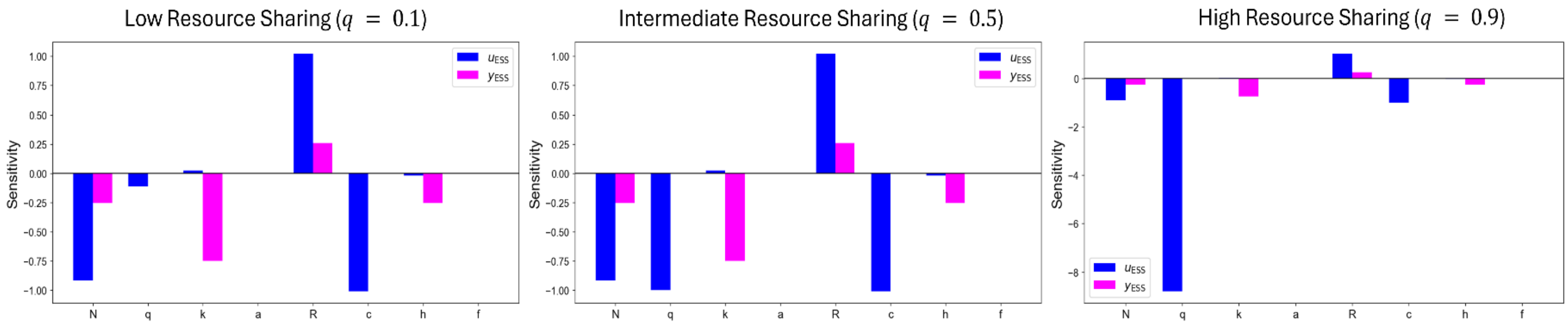


Fig. S7: Local normalized sensitivities of VEGF ($u$) and GLUT1 ($y$) expression at ESS at low, intermediate, and high resource sharing in a fixed neighborhood ($N = 10$).

##### 4.2 Dynamic Neighborhood

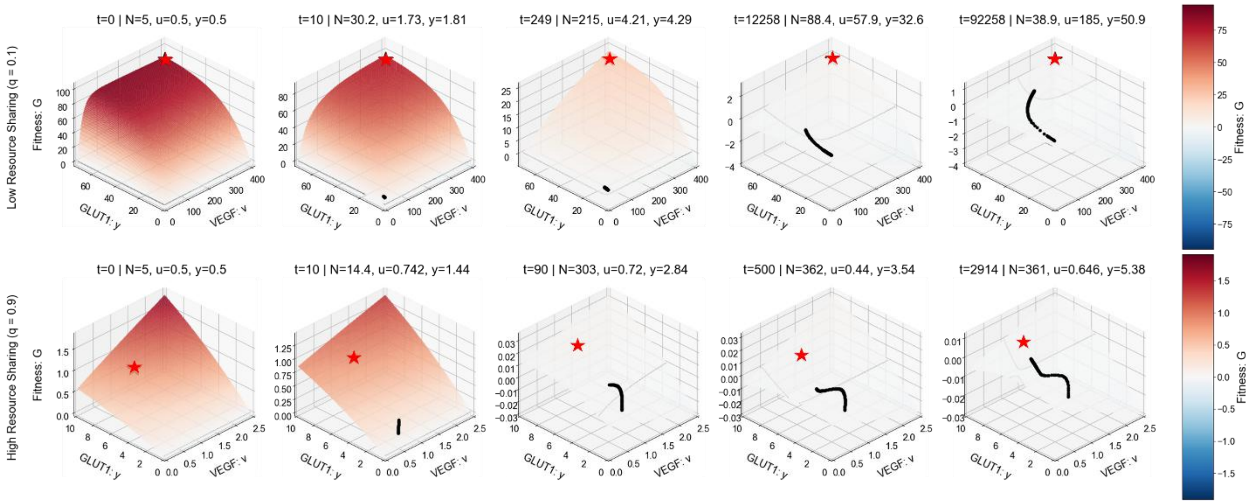


Fig. S8: Adaptive Landscape for Dynamic Neighborhood: Each graph represents a snapshot in time in the evolution of a focal cell introduced into the neighborhood. The population size ($N$), VEGF expression ($v$), GLUT1 expression ($y$) of the population at each time point is shown at the top of each graph (top = low resource sharing, bottom = high resource sharing). The black circles represent the trajectory of the population and the red star is the ESS. At equilibrium, the population reaches the red star and the fitness $G = 0$.

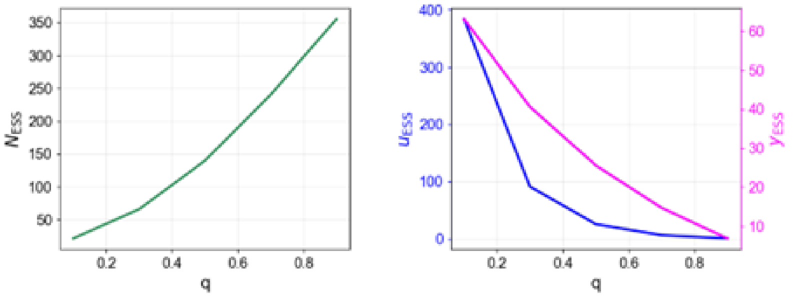


Fig. S9: Effect of the degree of resource sharing ($q$) on population size ($N$), VEGF expression ($u$), and GLUT1 expression ($y$) at ESS.

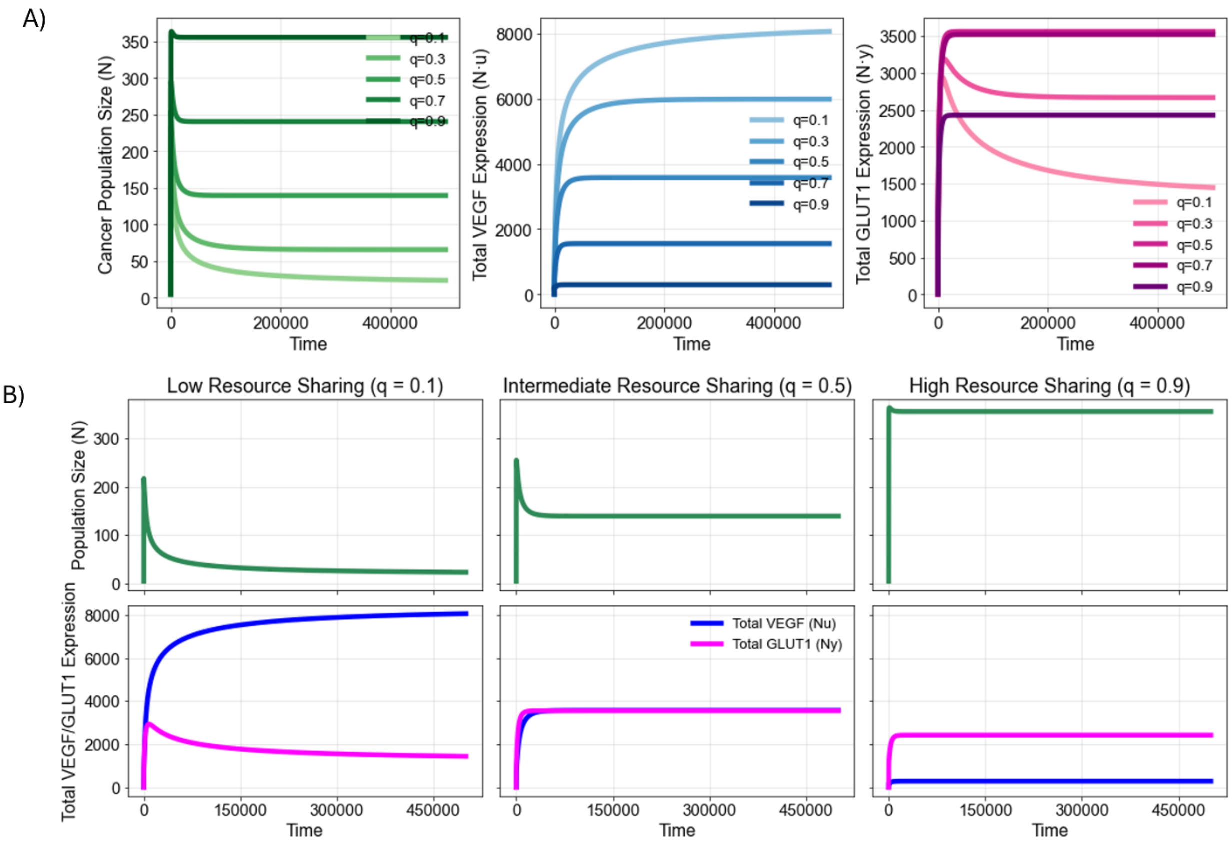


Fig. S10: A) Population Size ($N$), Total VEGF Expression ($Nu$), and total GLUT1 Expression ($Ny$) dynamics for different values of $q$. Darker the shade, higher the resource sharing ($q$). B) Comparison of ESS dynamics in low ($q = 0.1$), intermediate ($q = 0.5$), and high ($q = 0.9$) resource sharing neighborhoods.

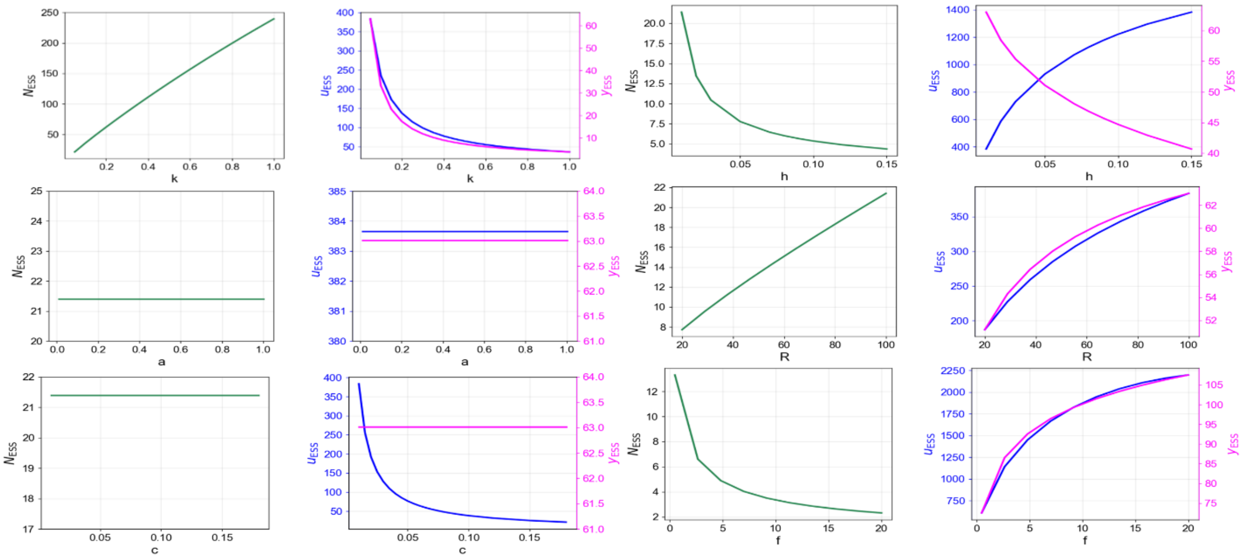


Fig. S11: Parameter sweeps for population size ($N$), VEGF expression ($u$), GLUT1 expression ($y$) at ESS in dynamic neighborhood for low resource sharing environment ($q = 0.1$).

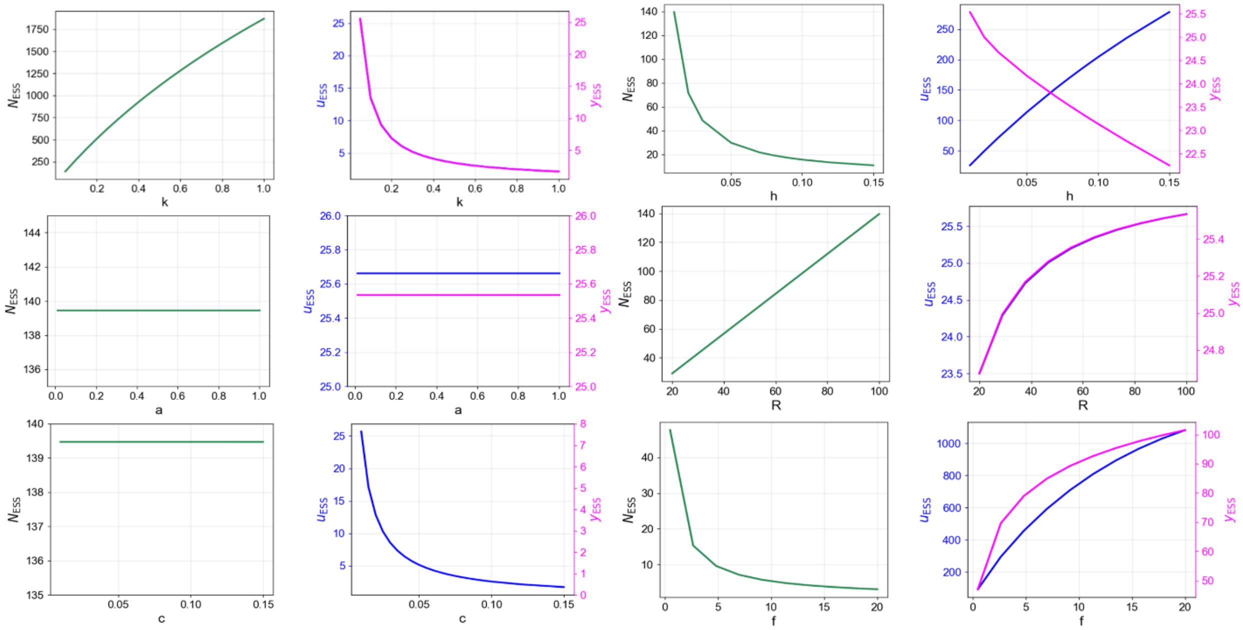


Fig. S12: Parameter sweeps for population size ($N$), VEGF expression ($u$), GLUT1 expression ($y$) at ESS in dynamic neighborhood for intermediate resource sharing environment ($q = 0.5$).

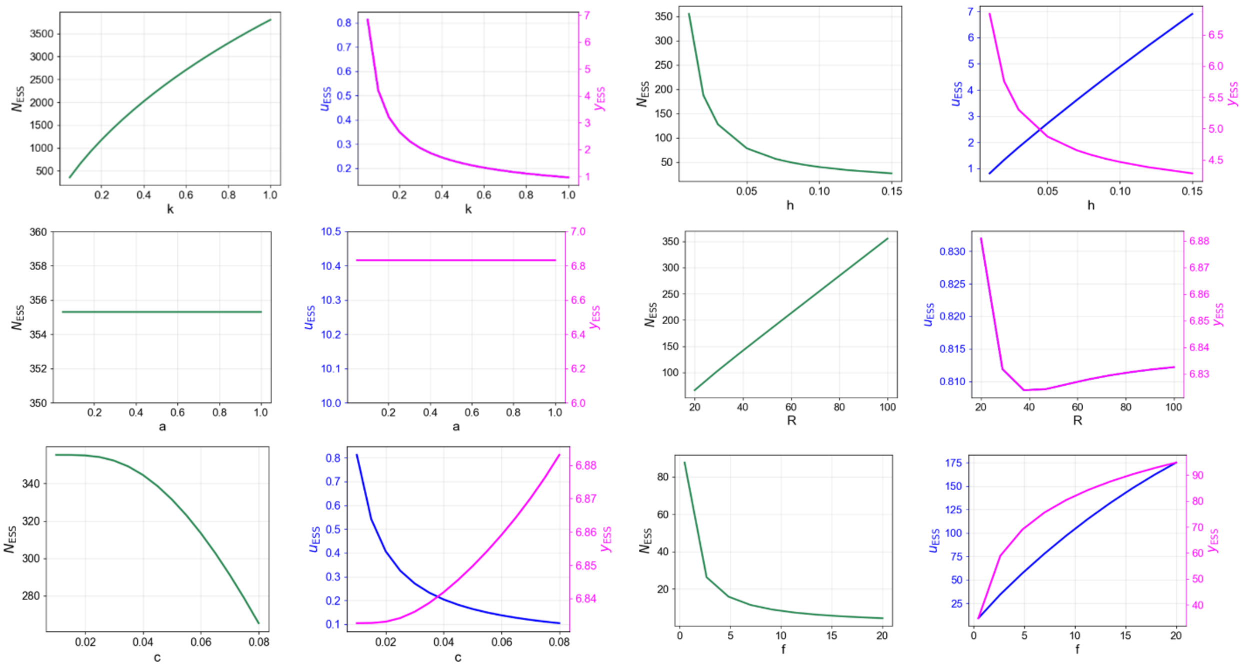


Fig. S13: Parameter sweeps for population size ($N$), VEGF expression ($u$), GLUT1 expression ($y$) at ESS in dynamic neighborhood for high resource sharing environment ($q = 0.9$).

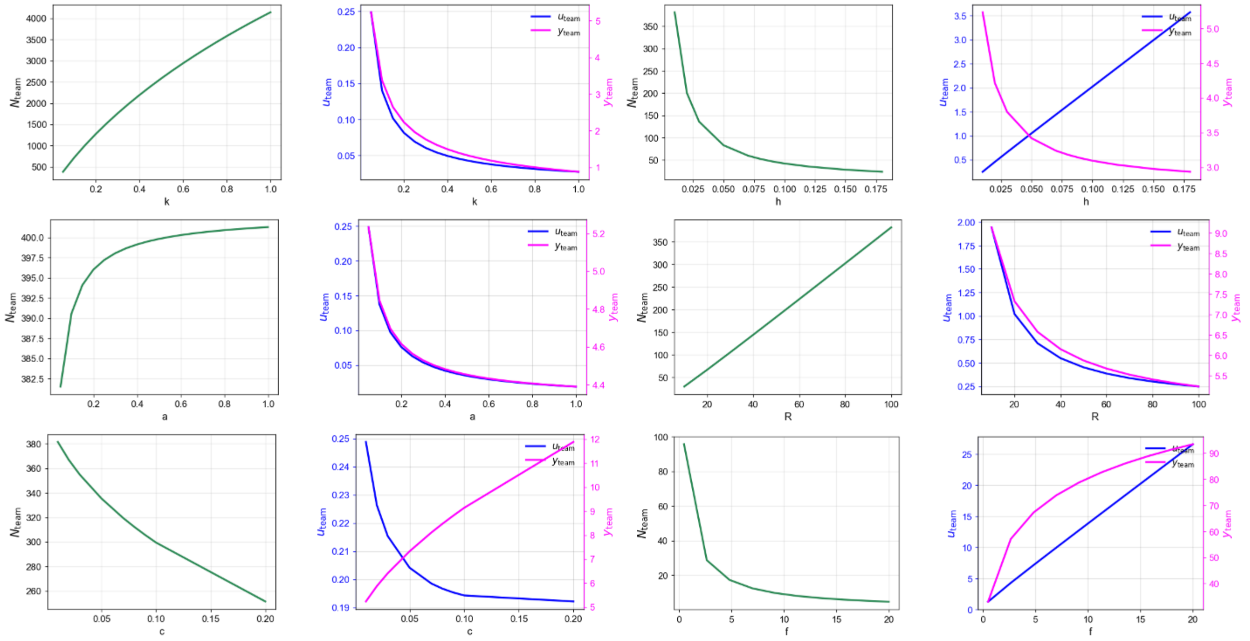


Fig. S14: Parameter sweeps for population size ($N$), VEGF expression ($u$), GLUT1 expression ($y$) at the team optimum in dynamic neighborhood.

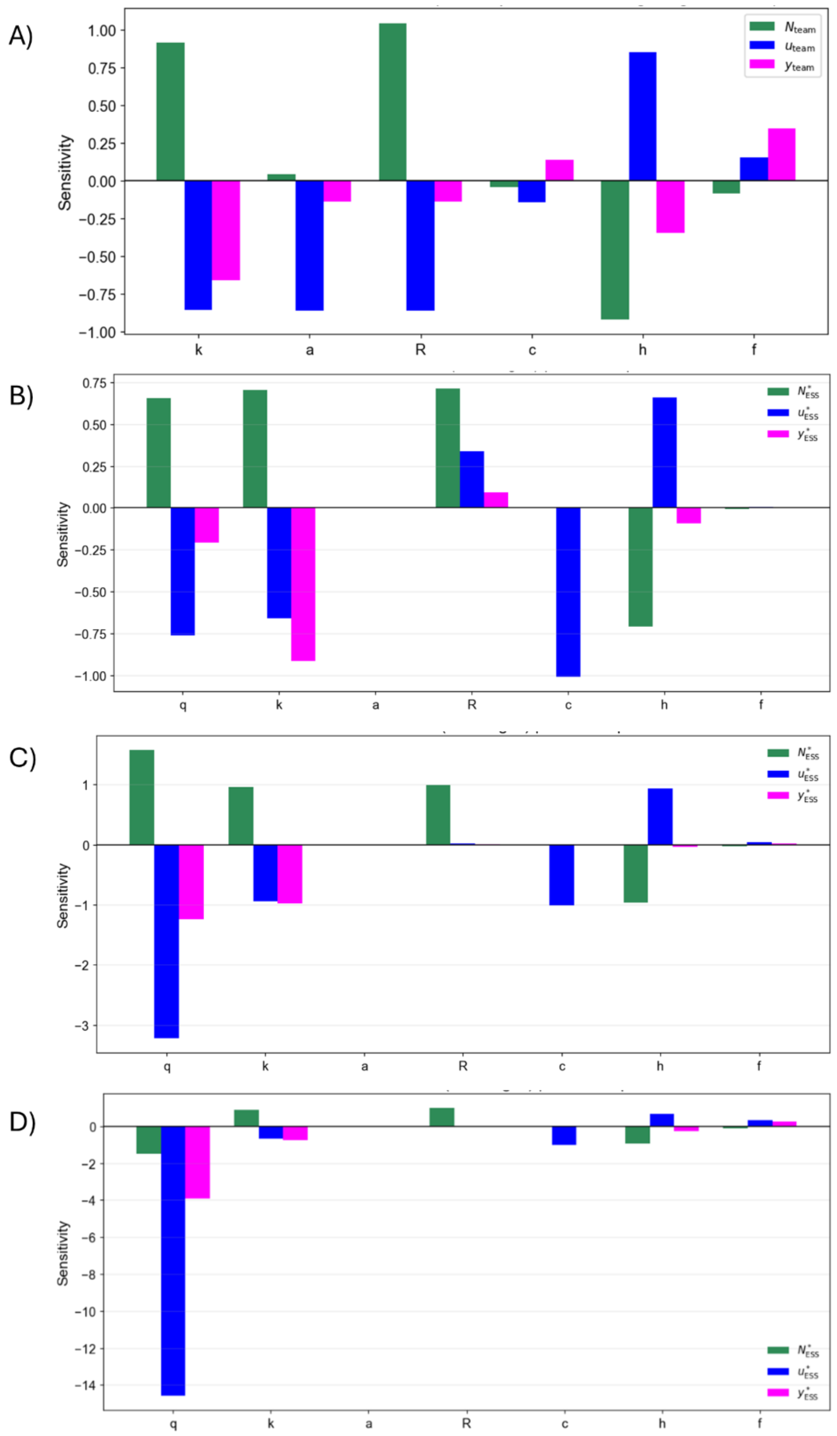


Fig. S15: Normalized local sensitivities for dynamic neighborhood: A) Team Optimum B) ESS for low resource sharing ($q = 0.1$) C) ESS for intermediate resource sharing ($q = 0.5$), and D) ESS for high resource sharing ($q = 0.9$).

#### 4.3 Therapy Dynamics

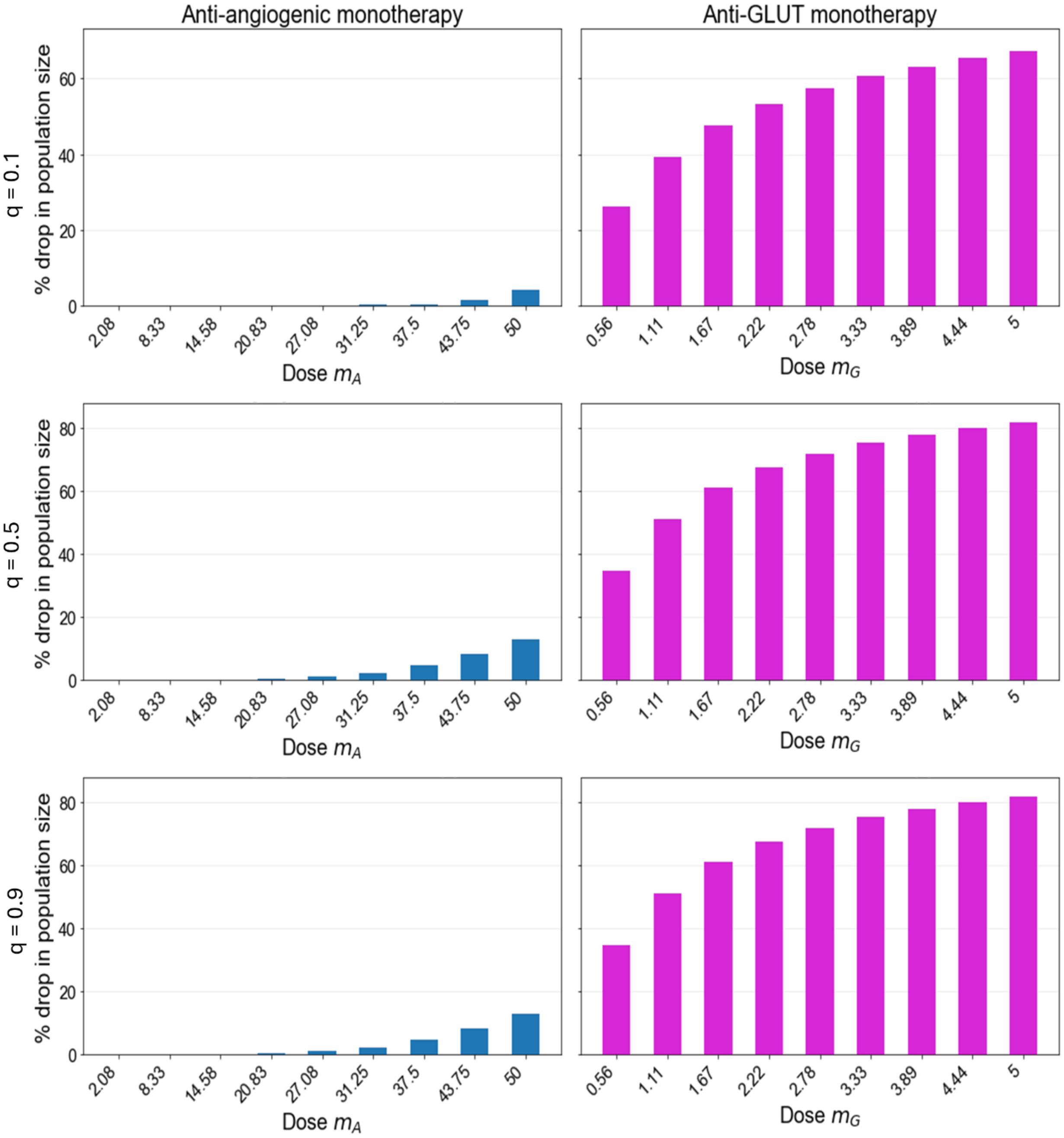


Fig. S16: Effect of different doses of monotherapies with anti-angiogenic and GLUT1 inhibitor on population size for low ($q = 0.1$), intermediate ($q = 0.5$), and high ($q = 0.9$) resource sharing.
